# RpiRc regulates RsbU to modulate eDNA-dependent biofilm formation and *in vivo* virulence of *Staphylococcus aureus* in a mouse model of catheter infection

**DOI:** 10.1101/783985

**Authors:** Adrien Fischer, Myriam Girard, Floriane Laumay, Anne-Kathrin Woischnig, Nina Khanna, Patrice François, Jacques Schrenzel

## Abstract

*Staphylococcus aureus* is a major human pathogen. Despite high incidence and morbidity, molecular mechanisms occurring during infection remain largely unknown. Under defined conditions, biofilm formation contributes to the severity of *S. aureus* related infections. Extracellular DNA (eDNA), a component of biofilm matrix released from apoptotic bacteria, is involved in biofilm structure and stability. In many bacterial biofilms, eDNA originates from cell lysis although eDNA can also be actively secreted or exported by bacterial membrane vesicles. By screening the Nebraska transposon library, we identified *rpiRc* as a biofilm regulator involved in eDNA regulation. RpiRc is a transcription factor from the pentose phosphate pathway (PPP) whose product is a polysaccharide intercellular adhesin (PIA) precursor. However, *rpiRc* mutant strain showed neither susceptibility to DispersinB® (a commercially available enzyme disrupting PIA biofilms) nor alteration of *ica* transcription (the operon regulating PIA production). Decreased biofilm formation was linked to Sln, an extracellular compound degrading eDNA in an autolysis independent pathway. Biofilm susceptibility to antibiotics in wt and mutant strains was tested using a similar protocol as the Calgary biofilm device. Involvement of RpiRc in *S. aureus* virulence was assessed *ex vivo* by internalization experiments into HEK293 cells and *in vivo* in a mouse model of subcutaneous catheter infection. While minimum inhibitory concentrations (MICs) of planktonic cells were not affected in the mutant strain, we observed increased biofilm susceptibility to almost all tested antibiotics, regardless of their mode of action. More importantly, the *rpiRc* mutant showed reduced virulence in both *ex vivo* and *in vivo* experiments related to decreased *fnbpA-B* transcription and eDNA production. RpiRc is an important regulator involved in eDNA degradation inside the matrix of mature PIA independent biofilms. These results illustrate that RpiRc contributes to increased antibiotic tolerance in mature bacterial biofilm and also to *S. aureus* cell adhesion and virulence during subcutaneous infection.

**Author summary:** Biofilm formation contributes to the severity of *Staphylococcus aureus* related infections. Biofilm matrix is mainly composed by polysaccharide intercellular adhesion (PIA), proteins and extracellular DNA (eDNA). By screening a mutant library of *S. aureus*, RpiRc was identified as a new regulator of eDNA dependent biofilm formation. How RpiRc regulates biofilm and its role in S. aureus virulence was studied in four different *S. aureus* strains. Deletion of RpiRc resulted in a pronounced decreased eDNA dependent biofilm formation, but not PIA dependent biofilm formation. Decreased biofilm formation was not related to increased autolysis, but was linked to extracellular compounds found in the supernatant of mutant biofilms. Sln was identified as one of this compound. RpiRc deletion also decreased biofilm recalcitrance (resistance) to selected antibiotics. Involvement of RpiRc in *S. aureus* pathogenesis was investigated *ex vivo* by internalization into HEK293 cells and *in vivo* in a mouse model of catheter infection. RpiRc deletion resulted in decreased virulence related to decreased expression of surface proteins like the fibronectin binding proteins A and B (FnbpA-B). These results illustrate that RpiRc contributes to increased antibiotic tolerance in mature bacterial biofilm and also to *S. aureus* cell adhesion and virulence during subcutaneous infection.

## Introduction

Biofilm is the most common mode of bacterial growth on medical devices and has also been reported on human tissues, *e.g.* during lung infection by *Pseudomonas aeruginosa*. Susceptible bacteria growing under biofilm conditions become recalcitrant to antibiotic, even in the presence of very high drug concentration (1, 2). Diagnostic and treatment of biofilm related infections remain an important challenge and the first guideline addressing this topic was recently published (3, 4).

Biofilms are sessile microbial communities attached to a surface and embedded in an extracellular matrix (5, 6). Biofilm formation is therefore the development of bacterial multilayers, encased within an exopolysaccharide glycocalyx (7), or any other extracellular matrix, mostly organized in a three dimensional structure. Formation of this structure consists of a primary attachment followed by bacterial accumulation, maturation and finally dissemination (8). An additional third step named exodus was recently added for *Staphylococcus aureus* biofilm regulated by nucleases (9). *Staphylococcus aureus* is a prevalent human pathogen able to form biofilms on indwelling devices (10–14), increasing the severity of several *S. aureus* related infections (*e.g.* cystic fibrosis, endocarditis, infectious arthritis, and catheter-related infections) (15–20).

In *S. aureus*, bacterial interactions with a surface or other cells are mediated by polysaccharide intercellular adhesin (PIA) (21), proteins like fibronectin binding proteins (FnBPs) (22–26), eDNA (27–31) or amyloid fibers composed by phenol soluble modulins (PSMs) (32, 33). PIA production in the biofilm matrix is regulated by the *ica* operon (*icaABCD*) (34). MRSA strains and some MSSA strains produce *ica*-independent biofilms that contain proteins and eDNA (25,26,29). These biofilms are commonly dispersed using proteases like SspB and ScpA or nucleases (35–37), respectively. USA300 biofilm formation requires FnBPs, but not other adhesins such as clumping factor A (ClfA) and protein A (Spa) (38).

eDNA is important for biofilm structuration and stability in almost all bacterial biofilms (39–42). It is also involved in bacterial attachment to the surface (41, 43) and in cell-to-cell interactions within the biofilm (44). In many bacteria, eDNA in the biofilm matrix originates from cell lysis, which is tightly regulated by the autolysins. In *S. aureus* two autolysis regulators are involved in this process: the major autolysin Atl (45) and the bacteriophage holin-antiholin like system Cid-Lrg (46), regulated by CidR and LytSR (47–49). Interestingly, studies reported active eDNA production using secretion vacuoles or other secretion mechanisms (44,50–53). We previously identified GdpS as a new actor in eDNA regulation (54). GdpS inhibits eDNA production by activating the eDNA inhibitor antiholin *lrgAB*.

Biofilm formation in *S. aureus* is tightly regulated by the quorum sensing (QS) system. *S. aureus* QS is under the control of the *agr* operon (55, 56). The *agr* system initiates the transcription of RNAIII, increasing extracellular proteases production and inhibiting cell adhesion to a surface. Proteins belonging to the microbial surface components recognizing adhesive matrix molecules (MSCRAMM), like the fibronectin binding proteins, are required for both bacterial cell adhesion to surface and interaction between bacteria and host cells (57). FnBPs mediate intercellular interaction and cell accumulation through low-affinity homophilic bonds during early phase of biofilm formation (38, 58), and are also required for biofilm maturation (45). RNAIII is an inhibitor of *fnbA* and *fnbB* transcription but *fnbA* is also under the control of SarA, another global regulator in *S. aureus*. SarA is under the control of the transcriptional regulator SigB (56, 59). SigB activity is tightly regulated by RsbU, leading to agr (RNAIII) inhibition and decreased extracellular proteases production, favoring FnBPs dependent biofilm formation (60, 61). The N3 sub-domain from the A domain of FnBPs is required for biofilm formation but not for fibronectin binding, meaning that binding to fibronectin and intercellular interaction during biofilm formation are independent (38, 62). Deletion of both genes is required for decreased biofilm formation and either *fnbA* or *fnbB* complementation alone restored MRSA biofilm amounts (38). SigB positively regulated *clfA* transcription while *clfB* transcription is enhanced by Rot, another transcription factor involved in *S. aureus* biofilm formation (63). CflB regulation by Rot is under the control of the agr system (64). ClfB like FnBPs are produced during early stages of biofilm formation and are involved in bacterial attachment to surface coated devices. ClfB is also required for *S. aureus* nasal colonization through direct interaction with nasal epithelial cells.

The Nebraska Transposon Library (NTL, (65)) was used to screen for genes involved in eDNA dependent biofilm formation. RpiRc was identified as an important *S. aureus* biofilm regulator and mechanisms under its control to modify eDNA and biofilm amounts were identified. The role of RpiRc in *S. aureus* interaction with host cells and virulence in a mouse model of catheter infection was also investigated and demonstrated RpiRc importance for *S. aureus in vivo* virulence

## Results

### Screening of the Nebraska Transposon Library to identify RpiRc

Starting from the 1920 mutant strains in the Nebraska Transposon Library (NTL), 108 genes involved in biofilm formation were identified (Figure S1). Experimental procedure focused on PIA and eDNA content testing addition of either DispersinB or DNaseI to the biofilm growth medium with these 108 mutant strains. For each experiment, several wt strains were added randomly in the 96 well plates to increase the relevance of the screening. Increased biofilm formation was observed in the center of the plate independently of the strains inoculated. To correct this bias all biofilms OD (Cristal Violet (CV)) were normalized twice with its row median and line median. This normalization resulted in a strong reduction in the difference observed between wt and mutant strains. For this reason, some genes previously described involved in the biofilm formation were missed but the probability to observe differences that are not true also decreased. From the 108 mutant strains identified, 16 showed modifications in PIA or eDNA content (Figure 1). This screening allowed the identification of RpiRc as an actor in *S. aureus* biofilm formation and eDNA regulation. Other targets already described involved in biofilm formation, eDNA or PIA production were identified like PknB, ClpP or SarA (Figure 1) (66). RpiRc presented the most promising results showing decreased biofilm formation correlated to the absence of DNaseI susceptibility, meaning absence of eDNA in *rpiRc* mutant biofilm. RpiRc is a transcription factor involved in the PPP pathway leading to the production of PIA precursor. The susceptibility to DispersinB was not modified for this mutant (Figure 1). USA300 JE2 is an MRSA and MRSA produce a Pbp2a which inhibits the production of PIA (67), meaning that any PIA regulation under the control of RpiRc could not be tested in this MRSA strain.

**Figure 1:**
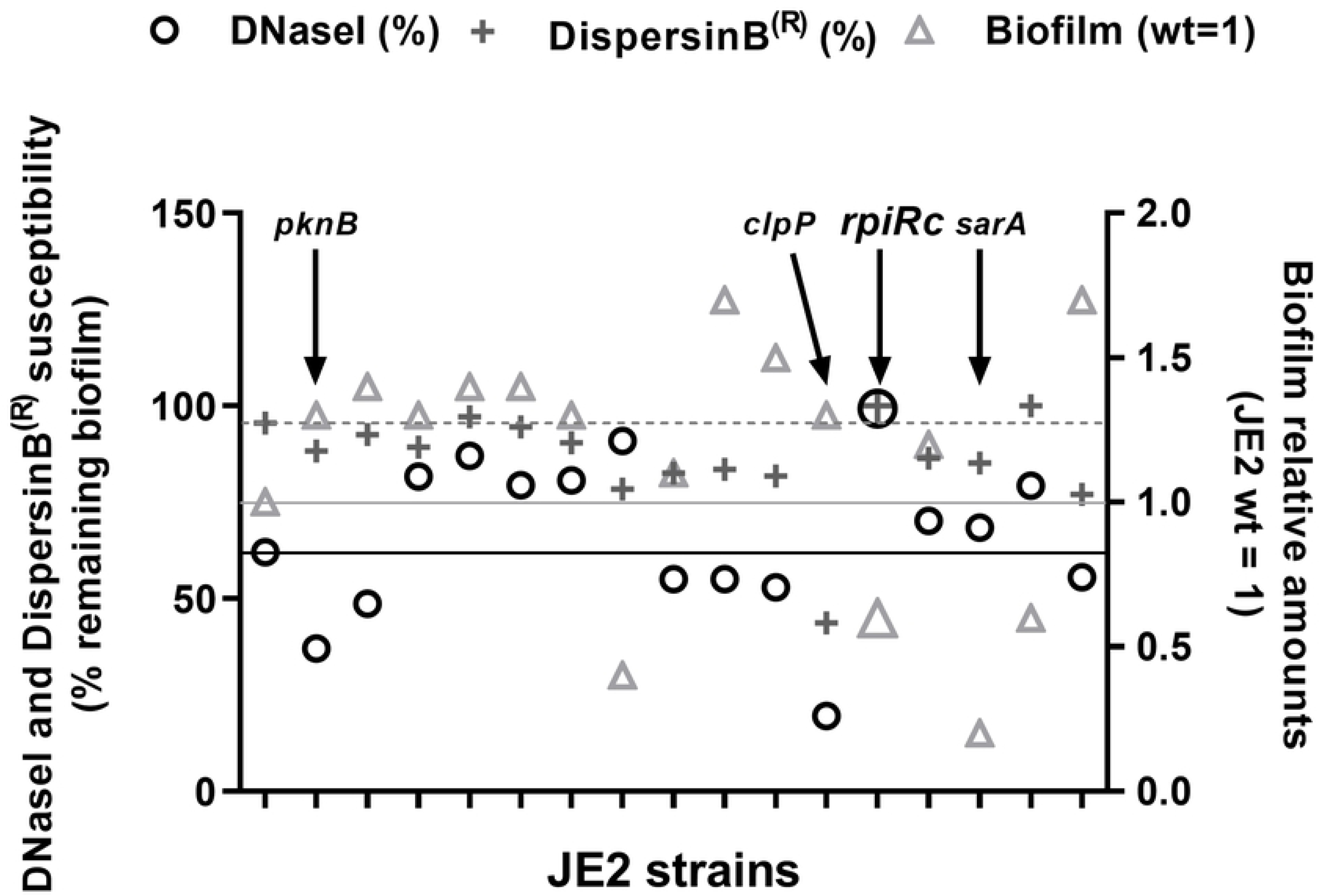
Mutants with altered biofilm formation and DNaseI or DisperinB susceptibility. Each column represents a mutant strain compared to the wt (lines). Biofilm amounts (triangles), DNaseI susceptibility (circles) and Dispersin B susceptibility (cross) are represented vertically for each. Lines represent USA300 JE2 wt relative amount of biofilm (continuous gray line) and remaining biofilm after DNaseI (continuous black line) or DispersinB treatment (discontinuous gray line). Some known targets are indicated on the graph like *pknB*, *clpP* or *sarA*. The selected target for this study, *rpiRc* is also shown with bigger sized symbols.

### RpiRc regulates eDNA dependent mature biofilm of different strains

To confirm *rpiRc* involvement in biofilm formation and eDNA production or stability, *rpiRc* mutation was back transduced in USA300 JE2 wt. Biofilm was grown in 24 well plates following usual protocol for biofilm and eDNA quantification from the laboratory. Three time points were tested, 1.5, 6 and 24 hours, to determine in which phase of biofilm formation RpiRc could be involved. *rpiRc* only altered mature biofilm formation (Figure 2A), corresponding to an important decreased eDNA production (Figure 2B). *rpiRc* also alters eDNA production in 6 hours old biofilms, without affecting biofilm amounts. RpiRc complementation (JE2 Δ*rpiRc* Nat7) restored biofilm formation and eDNA amounts (Figure 2A-B).

**Figure 2:**
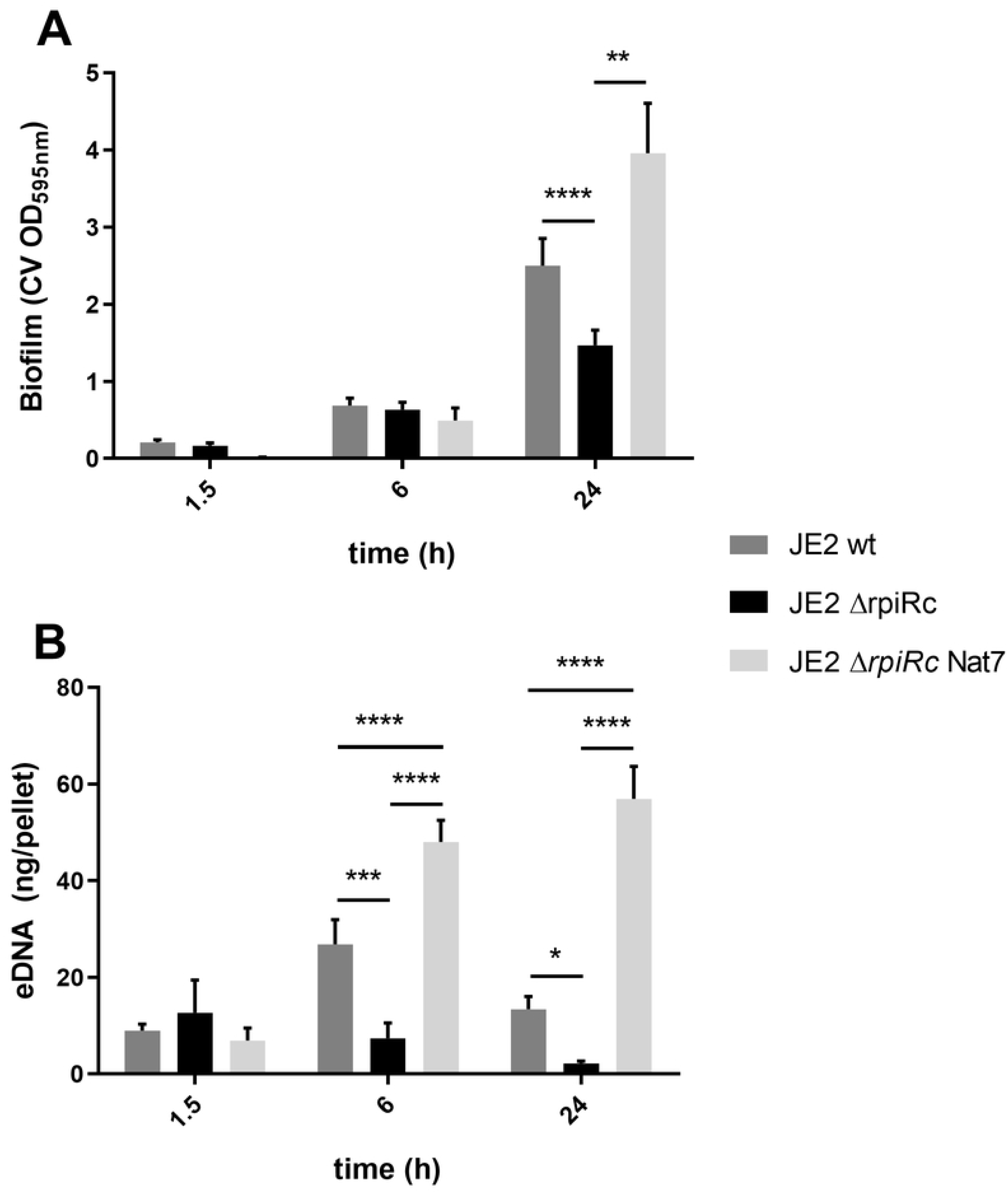
RpiRc regulates biofilm formation and eDNA in USA300 JE2. USA300 JE2 wt, Δ*rpiRc* (re-transduced) and complementation (Nat7) were tested for (A) biofilm formation using cristal violet staining at different time points (1.5, 6 and 24h) and, (B) eDNA quantification at different time points (1.5, 6 and 24h). Data represents mean of 10 independent experiments; *; **; ***; ****: p<0.05; <0.01; <0.001; <0.0001 respectively.

RpiRc was then deleted in three other strains, two clinical isolates UAMS-1 and SA564 and one laboratory strain producing PIA, SA113 (Figure 3). All mutant strains had similar growth curves compared to their wt counterpart (Figure S2). Mature biofilm formation for UAMS-1 and SA564 was affected by *rpiRc* deletion (Figure 3 A-B). Six hours old biofilm were not affected by *rpiRc* deletion in UAMS-1, while affected in SA564 to a similar extend than mature biofilm (Figure 3B). Strains UAMS-1 Δ*rpiRc* and SA564 Δ*rpiRc* produced less eDNA at 6h and 24h compared to wt, SA564 Δ*rpiRc* showing a complete abolition of eDNA amounts in the biofilm. RpiRc complementation always restored both biofilm formation and eDNA production (Figure 3 A-B-D-E). The three clinical strains tested here showed similar role for *rpiRc* as a mature biofilm formation and eDNA regulator. This is not true with the laboratory strain SA113. Deletion of *rpiRc* did not alter biofilm formation nor eDNA amounts in the biofilm (Figure 3 C-F). RpiRc complementation resulted in a decreased biofilm formation at 6h for SA113 using our standard CV staining assay. This observation could be correlated to delayed growth rate observed for all complemented strains (Figure S2 A-B-C-D; 5 to 12 hours delay, SA113 Nat7 showing the higher delayed growth). Therefore it was hypothesized that SA113 Nat7 produced less biofilm in our experimental conditions due to its slow growth. To test this hypothesis, and reject a probable contrary role of RpiRc in SA113, the biofilm index of SA113 wt, mutant and complemented strain was estimated (Figure S3) referring to previously published methods (68–71). Biofilm index for SA113 Nat7 was similar to wt and *rpiRc* mutant, confirming that decreased OD_595nm_ of CV was a consequence of Nat7 slow growth and that RpiRc has no effects on biofilm formation and eDNA production in the laboratory strain SA113. This suggests again that RpiRc does not regulate PIA dependent biofilm formation.

**Figure 3:**
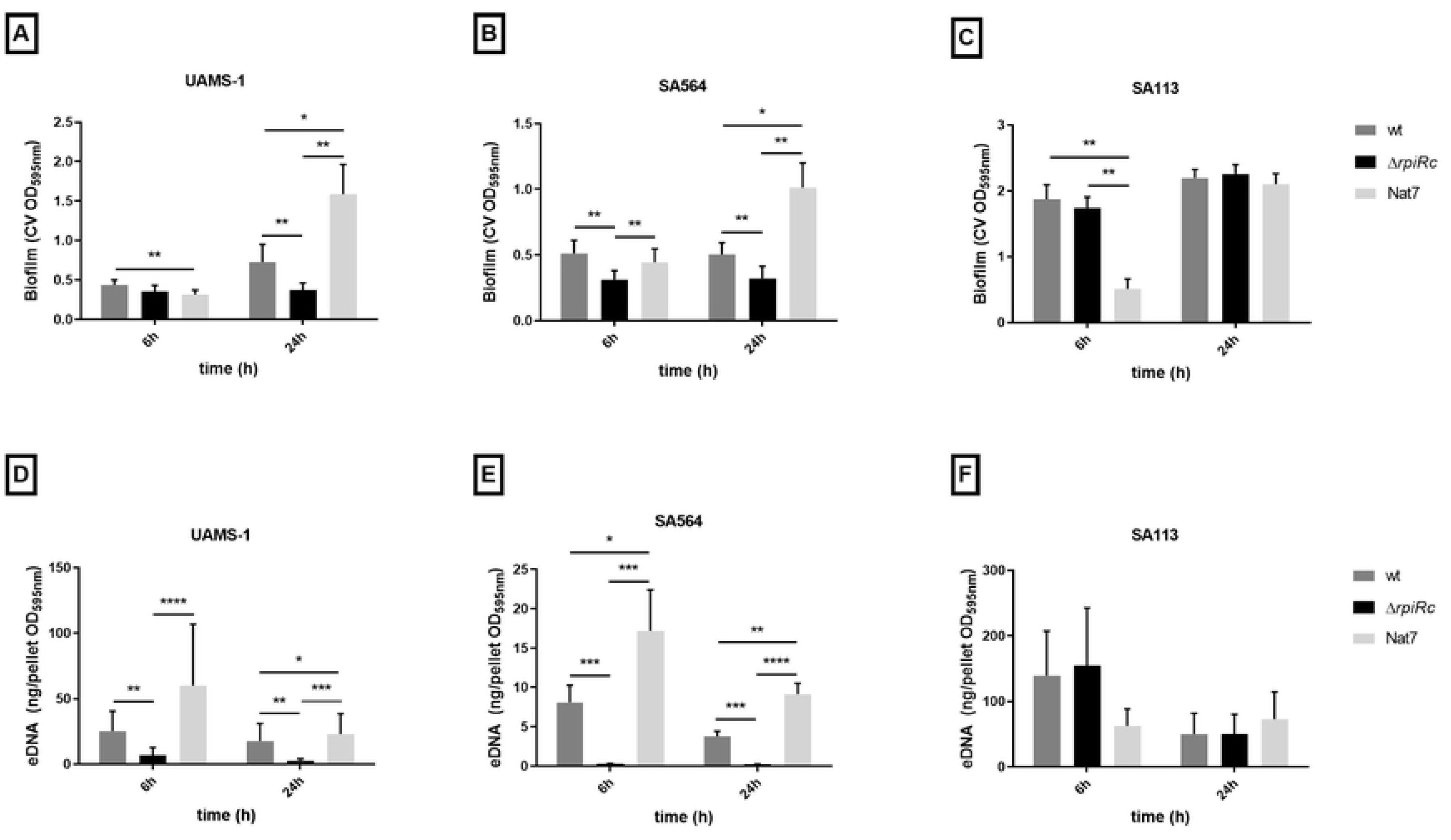
RpiRc regulates biofilm formation and eDNA in other backgrounds. wt, Δ*rpiRc* and complementation (Nat7) were tested for biofilm formation in UAMS-1 (A), SA564 (B) and SA113 (C) at different time points (6h and 24h). eDNA amounts were also quantified in the wt, Δ*rpiRc* and complementation of UAMS-1 (D), SA564 (E) and SA113 (F) after 6h and 24h of biofilm formation. Data represents mean of 5 independent experiments; *; **; ***; ****: p<0.05; <0.01; <0.001; <0.0001 respectively.

Compared to wt, *rpiRc* complementation in the clinical strains always showed increased mature biofilm formation (t=24h) and eDNA production (t=6h and 24h) suggesting a dose dependent role of RpiRc in biofilm formation, the more RpiRc, the more biofilm and eDNA are produced (Figure 3). Again this is not true for SA113 (Figure 3 C-F).

### RpiRc regulates RsbU to control eDNA amounts on mature biofilms

The regulation of eDNA amount in the biofilm of *rpiRc* mutant strains is not related to increased induced autolysis (Figure S4). Oxacillin induced autolysis also did not show differences for USA300 JE2 strain (Figure S5), therefore suggesting another mechanism for eDNA regulation by RpiRc. SA113 was not affected by *rpiRc* deletion neither for eDNA nor biofilm amounts. SA113 is clonal to NCTC8325 and carries mutations in important *S. aureus* regulators like TcaR or RsbU. RsbU is required for *sigB* activity, regulating extracellular proteases production as well as cell wall metabolism or biofilm formation. It was hypothesized that absence of modification in SA113 Δ*rpiRc* is related to *rsbU* mutation. The transcription level of *rsbU* in wt and *rpiRc* mutant USA300 JE2 strains under biofilm growth conditions was determined and showed a non-statistically significant increase in the mutant strains in 3h old biofilms (Table 1 and Tables S1 and S2), meaning that *rpiRc* may regulate biofilm formation and eDNA through RsbU regulation. To confirm this hypothesis, *rsbU* was deleted in UAMS-1 and SA564, wt and Δ*rpiRc*. Biofilm formation and eDNA amounts were measured in these strains (Figure 4). RsbU single deletion was responsible for moderate increased biofilm formation in SA564 at 6h and 24h (Figure 4B) as well as increased eDNA amounts at 6h and 24h for SA564 (Figure 4D) and 24h for UAMS-1 (Figure 4C). *rsbU* deletion in *rpiRc* mutants resulted in a low but significant biofilm formation increase at 6h for UAMS-1 (Figure 4A) and at 24h for both strains (Figure 4A-B). Double mutants showed at both time points with UAMS-1 and SA564 a complete restoration of eDNA amounts compared to each wt strains (Figure 4C-D). eDNA regulation in growing and mature biofilms by RpiRc is mediated by RsbU.

**Figure 4:**
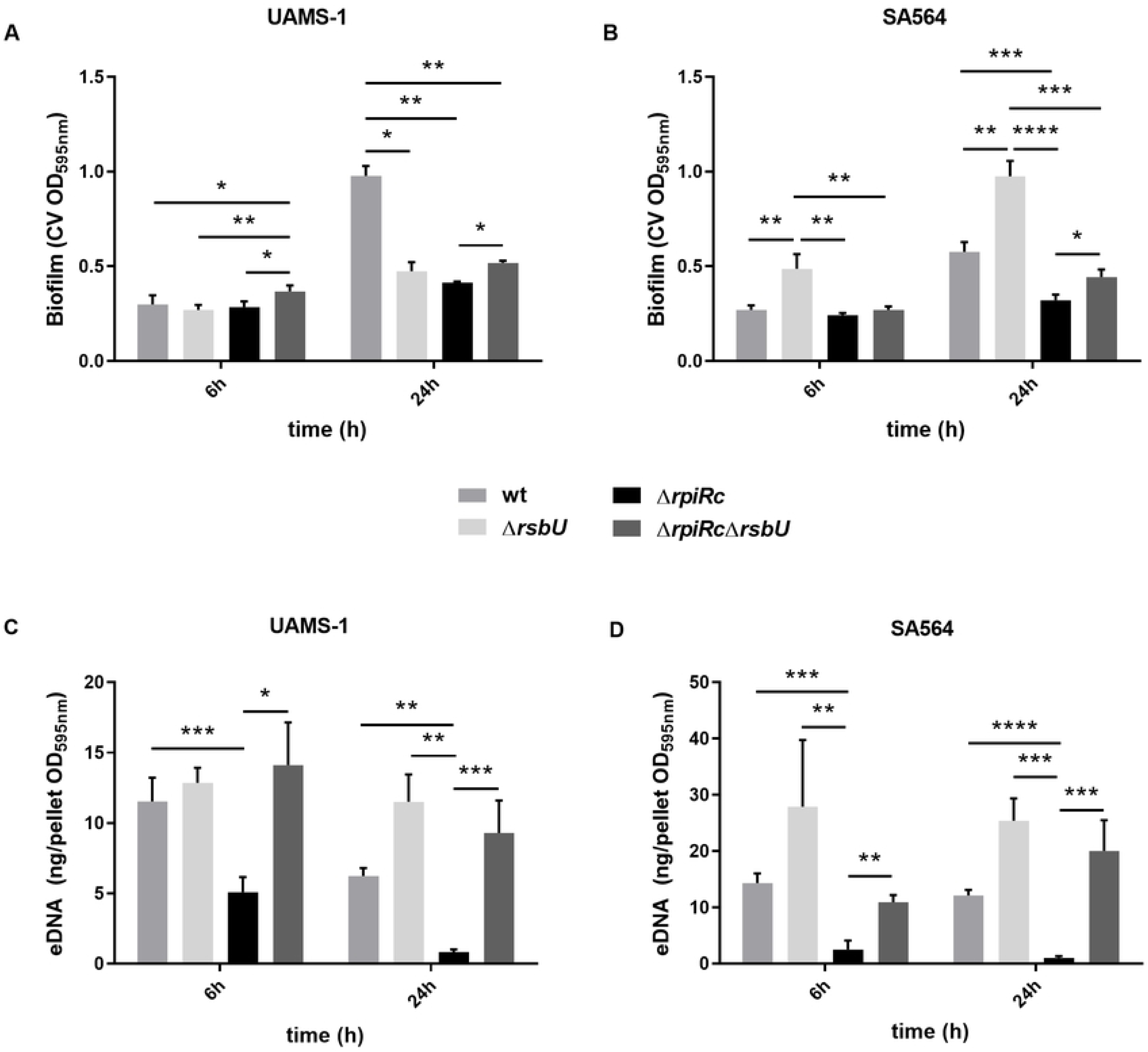
RpiRc regulates RsbU to control eDNA amounts In biofilms. wt, Δ*rsbU*, Δ*rpiRc* and double mutant Δ*rsbU*Δ*rpiRc* were tested for biofilm formation in UAMS-1 (A) and SA564 (B) and eDNA amounts were quantified in UAMS-1 (C) and SA564 (D) after 6h and 24h of biofilm formation. Data represents mean (+ standard error of the mean) of 3 independent experiments; *; **; ***; ****: p<0.05; <0.01; <0.001; <0.0001 respectively.

**Table 1:** mRNA relative amounts in 3h and 6h old biofilms.

| mRNA<br>biofilm | *rsbU* |  | *agrA* |  | *RNAIII* |  |
| --- | --- | --- | --- | --- | --- | --- |
| | USA300<br>JE2 $\Delta rpiRc$ | USA300 JE2<br>Nat7 | USA300<br>JE2 $\Delta rpiRc$ | USA300 JE2<br>Nat7 | USA300<br>JE2 $\Delta rpiRc$ | USA300 JE2<br>Nat7 |
| 3h | 1.30 | **0.70** | 2.00 | **0.14** | **10.0** | **0.09** |
| 6h | 0.90 | 0.70 | **1.30** | **0.14** | **2.80** | **0.01** |

qRT-PCR was performed on USA300 JE2 wt, *rpiRc* mutant and complemented strains. Measures were repeated on six independent biological replicates. USA300 JE2 wt strain was used as reference (relative amount=1) for both time points. Bold values are statistically different from wt ( $p < 0.05$ , paired ttest). Standard deviation and ttest results are presented in Tables S1 and S2.

### RpiRc deletion promotes eDNA dispersion and degradation

RpiRc regulates the amount of eDNA on growing and mature biofilms but only affects mature biofilm amounts. It was aimed at deciphering if RpiRc involves eDNA production/release regulation or stability/degradation on the matrix. If RpiRc is involved in eDNA production, adding eDNA to the biofilm should restore wt eDNA amounts in the mutant biofilm. eDNA extracted (eeDNA for exogenous eDNA) from a strain containing an erythromycin cassette (*ermC*) was added to culture medium and specific primers for this cassette were designed. This specific primer pair did not amplify eDNA from USA300 JE2 wt or Δ*rpiRc* in absence of exogenous *ermC*. The amount of eeDNA incorporated in the wt and mutant biofilm were measured at four different time points (Figure 5A; 1.5, 3, 6 and 24h). eeDNA incorporation in the biofilm of both wt and mutant biofilms was measured. No differences of eeDNA amounts could be observed between wt and *rpiRc* mutant in early and growing biofilms (1.5, 3 and 6h). Mature biofilm of (24h) USA300 JE2 Δ*rpiRc* contained significantly less eeDNA compared to wt. This suggests that either *rpiRc* mutant strain no longer binds eDNA or that eDNA is degraded. eeDNA in the supernatant was measured for each time point (Figure 5B). No differences between wt and mutant strain were observed, meaning that decreased eeDNA in USA300 JE2 Δ*rpiRc* mature biofilm is most certainly related to increased degradation compared to the wt strain. To confirm this hypothesis, supernatant of USA300 JE2 Δ*rpiRc* from mature biofilm was added to wt mature biofilm and eDNA amounts were measured at different time points after SN addition (Figure 5C; 0.25, 1, 6 and 24h). Addition of wt mature biofilm supernatant on wt mature biofilm (wt + wt SN) was used as supernatant modification control. This condition always showed increased eDNA compared to the control without medium modification (wt ctl). When SN of USA300 JE2 Δ*rpiRc* mature biofilm was added to wt mature biofilm, eDNA amounts rapidly decreased compared to addition of wt mature biofilm SN. eDNA amounts in the wt biofilm in presence of USA300 JE2 Δ*rpiRc* SN were statistically lower after 15 minutes, 1 hour and 6 hours. After 24h, the difference is no longer statistically significant, but remains lower. The SN of *rpiRc* mutant biofilm contains a protein (enzyme) or a metabolite degrading eDNA in the matrix. As previously demonstrated, eDNA is involved in biofilm recalcitrance to vancomycin (72). Deletion of RpiRc in strain JE2 resulted in decreased biofilm recalcitrance for vancomycin, oxacillin, gentamycin, rifampicin and ciprofloxacin compared to the wt strain (Figure S6). However no difference using e-test could be observed with any of these antibiotics (Table S3).

**Figure 5:**
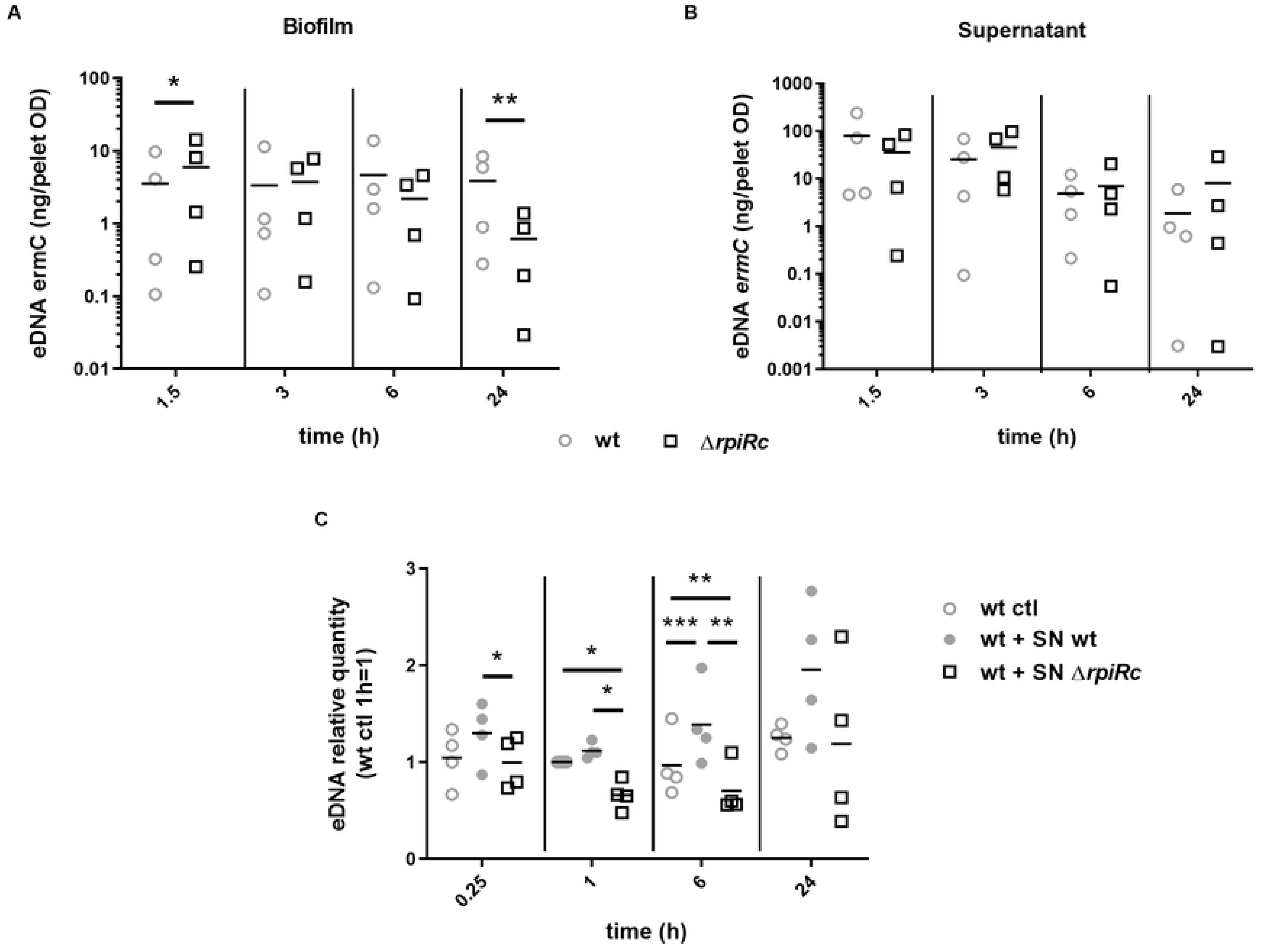
RpiRc promotes eDNA dispersion and degradation. eDNA containing an erythromycin resistance cassette was added to culture medium at the beginning of biofilm growth and the amount of this eeDNA in the biofilm matrix (A) of USA300 JE2 wt and mutant strain as well as in their supernatant (B) after 1.5h, 3h, 6h and 24h of biofilm growth. (C) Supernatant of 24h old USA300 JE2 Δ*rpiRc* strain (wt + SN Δ*rpiRc*) or USA300 JE2 wt strain (wt + SN wt) were added to a 24h old wt biofilm and eDNA quantified 15minutes, 1h, 6h and 24h after SN addition. As a control, eDNA was quantified from wt biofilm without supernatant modification (wt ctl). Data represents 4 independent experiments; *; **; ***: p<0.05; <0.01; <0.001 respectively.

RpiRc regulates eDNA stability on growing and mature biofilms most probably by the regulation of an extracellular nuclease. To test this hypothesis, both *S. aureus* extracellular nucleases *nuc1* and *nuc2* were deleted in SA564 wt and SA564 Δ*rpiRc* strains (Figure S7 and Table S4). Deletion of either *nuc1* or *nuc2* did not modify *rpiRc* dependent eDNA decrease in mature biofilm nor restored biofilm formation. RpiRc regulates an unknown secreted factor degrading eDNA in the matrix of *S. aureus* biofilm that remains to be identified. Supernatants of mature USA300 JE2 wt and Δ*rpiRc* biofilms were purified and proteins separated on SDS acrylamide gel. Mutant supernatant showed the highest intensity protein band at 35KDa while wt supernatant only presented a low intensity band (Figure S8). The composition of the 35KDa band was analyzed using LC-MS. A total of 90 different proteins were identified (Table S5). As expected a large number of proteins were over-produced in the mutant supernatant, representing mainly virulence determinant like the PVL, hemolysins (alpha, beta and gamma), the serine and cysteine proteases SspA and SspB (under the control of RsbU/SigB), the foldase protein PrsA (down-regulated) and other surface proteins like MecA, Sbi (down-regulated) or the major autolysin Atl (Table 2). Some proteins identified did not match the selected proteins weight analyzed (e.g around 35KDa). This is the case for SspB, Sbi, Atl and MecA. It is not clear whether these proteins are more degraded or if they are over-produced (more peptides form normal degradation found at lower weight). Three nucleases were also identified in the supernatants, Nuc1, Sln and AdsA (Table 2). Nuc1, which was already tested by deletion and did not reveal involved in *rpiRc* regulatory pathway, was over-produced in the wt supernatant, confirming the deletion results (Figure S7) and Nuc1 independent decreased eDNA amounts. Sln and AdsA are over-produced in the mutant supernatant, suggesting one or both as potential targets for RpiRc dependent eDNA regulation and biofilm formation. Sln had a weight of 33KDa while AdsA had a weight of 83KDa (Table 2). For this reason, Sln was selected for further mutagenesis experiments.

**Table 2:** Selected proteins identified in supernatants of wt and Δ*rpiRc* mature biofilms.

| Identified Proteins | Accession Number | MW (KDa) | gene name | $\Delta rpiRc/wt$ |
| --- | --- | --- | --- | --- |
| Thermonuclease | A0A0E1VLD0_STAA3 | 24 | *nuc1* | 0<sup>ψ</sup> |
| 5'-nucleotidase, lipoprotein e(P4) family | A0A0E1VNB4_STAA3 | 33 | *sln* | 2* |
| HPr kinase/phosphorylase | A0A0E1VLJ4_STAA3 | 34 | *hprK* | 13* |
| Beta-channel forming cytolysin | A0A0E1VG99_STAA3 | 35 | *hlb* | 15* |
| Panton-Valentine leukocidin, LukS-PV | A0A0H2XI46_STAA3 | 35 | *lukS-PV* | 1.8 |
| Foldase protein PrsA | A0A0E1VIE3_STAA3 | 36 | *prsA* | 0.2 |
| Alpha-hemolysin | A0A0H2XEW5_STAA3 | 36 | *hla* | 3.0 |
| Serine protease | A0A0H2XFC3_STAA3 | 36 | *sspA* | 2.9 |
| Leukotoxin LukD | A0A0H2XFN4_STAA3 | 37 | *lukD* | 23* |
| Gamma-hemolysin component B | A0A0E1VN13_STAA3 | 37 | *hlgB* | 2.5 |
| Panton-Valentine leukocidin, LukF-PV | A0A0H2XFK0_STAA3 | 37 | *lukF-PV* | 3.6 |
| Cysteine protease | A0A0H2XGH9_STAA3 | 45 | *sspB* | 2.3 |
| Immunoglobulin-binding protein Sbi | SBI_STAA3 | 50 | *sbi* | 0.0 |
| Penicillin-binding protein 2 | A0A0H2XHY4_STAA3 | 76 | *mecA* | 16* |
| 5'-nucleotidase family protein | A0A0H2XI02_STAA3 | 83 | *adsA* | 3* |
| Autolysin | A0A0H2XIJ3_STAA3 | 137 | *atl* | 3.2 |

$\Delta rpiRc/wt$ represents the ratio of the sum of peptide counts from the duplicate for $\Delta rpiRc$ and wt
\* no peptide identified in wt supernatant for this protein
<sup>ψ</sup> no peptide identified in $\Delta rpiRc$ supernatant for this protein

Sln corresponds to a 5’-nucleotidase, lipoprotein. This protein carries a domain to degrade riboside from the extracellular medium. This is an interesting target as *rpiRc* deletion should induce a decreased PPP dependent nucleotide synthesis. This gene was named *sln* for Staphylococcal lipoprotein nucleotidase. Sln was deleted in SA564 wt and Δ*rpiRc*, and biofilm formation and eDNA amounts were estimated at different time points of biofilm formation (3h, 6h and 24h). No differences in biofilm amounts could be observed at any stage of biofilm growth between SA564 wt and Δ*sln* as well as between SA564 Δ*rpiRc* and Δ*rpiRc*Δ*sln* (Figure 6-A). SA564 Δ*rpiRc* and Δ*rpiRc*Δ*sln* produce less biofilm compared to SA564 wt and Δ*sln* strains at any stage of biofilm growth (Figure 6-A). A small decrease in eDNA amounts could be monitored in the double mutant Δ*rpiRc*Δ*sln* compared to single mutant SA564 Δ*rpiRc* at 3h and 24h (0.8 and 0.7 fold change respectively, Figure 6-B). After 6h of biofilm formation, SA564 Δ*rpiRc*Δ*sln* showed increased eDNA amounts compared to SA564 Δ*rpiRc* (1.7 fold change, Figure 6-B). The amount of eDNA recovered from SA564 Δ*rpiRc* between 6h and 24h is not statistically different (p=0.084), while different for SA564 Δ*rpiRc*Δ*sln* showing a 3.8 fold decrease at 24h compared to 6h (p=0.0005), strongly suggesting that Sln is involved in RpiRc dependent eDNA regulation. Deletion of *sln* in SA564 Δ*rpiRc* strain do not restore biofilm amount neither completely restored eDNA amounts meaning that other regulator than Sln must be involved in RpiRc dependent biofilm formation and eDNA regulation pathways. Another hypothesis is the regulation of a protease degrading surface proteins involved in the binding of eDNA, unbound eDNA being more susceptible to extracellular DNases than bound eDNA.

**Figure 6:**
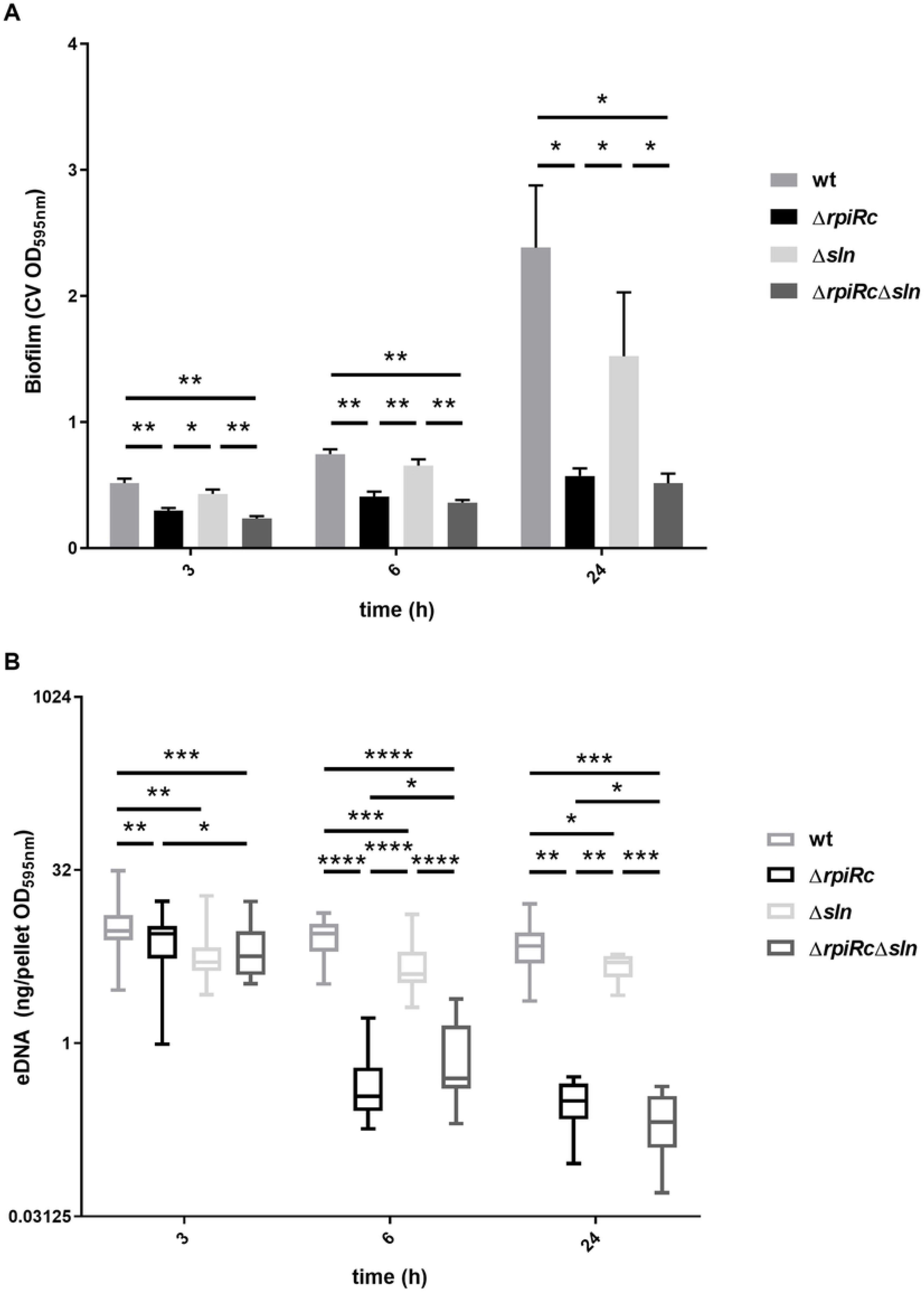
Sln is part of the RpiRc dependent eDNA regulatory network. wt, Δ*rpiRc*, Δ*sln* and double mutant Δ*rpiRc*Δ*sln* were tested for biofilm formation in strain SA564 (A) and eDNA amounts were quantified (B) after 3h, 6h and 24h of biofilm formation. Data represents mean of 6 (for biofilm formation) and interleaved box and whiskers with 5-95 percentile of 16 (for eDNA) independent experiments. For eDNA quantification a paired non-parametric Wilcoxon statistical test was performed; *; **; ***; ****: p<0.05; <0.01; <0.001; <0.0001 respectively.

### RpiRc is involved in the regulation of MSCRAMMs

Surface proteins belonging to the MSCRAMMs are important structural and adhesive component for *S. aureus* pathogenesis and biofilm formation in PIA independent biofilm forming strains like MRSA (USA300), UAMS-1 or SA564. To elucidate their role in RpiRc dependent biofilm formation, RNA from 3h and 6h old biofilms were extracted and the expression levels of *fnbA-B*, *clfA-B* and *fbp* were compared between USA300 JE2wt, Δ*rpiRc* and Nat7. After 3h of biofilm growth, expression levels of all these genes were altered in the mutant strain, *fnbA-B* and *clfB* showing decreased expression while *clfA* and *fbp* were two fold more expressed (Figure 7A). Complementation always restored the wt phenotype. Over-expression of *rpiRc* resulted in an important increased expression of *fnbA* and *fnbB*. After 6 hours of biofilm growth, *fnbA* and *B* remained down-regulated in the mutant strain (Figure 7B). The mRNA amounts of *clfA*, *clfB* and *fbp* were similar between the wt and the mutant strain. Again, the complementation presented huge increased *fnbA* and *B* expression (Figure 7B). In addition, a significant decreased *clfA* expression and increased *clfB* expression was observed (Figure 7B, Nat7) compared to the transcription profile at 3hours of biofilm formation (Figure 7A). RpiRc is involved in the transcription regulation of MSCRAMMs genes with a particular emphasis for *fnbA-B* genes.

**Figure 7:**
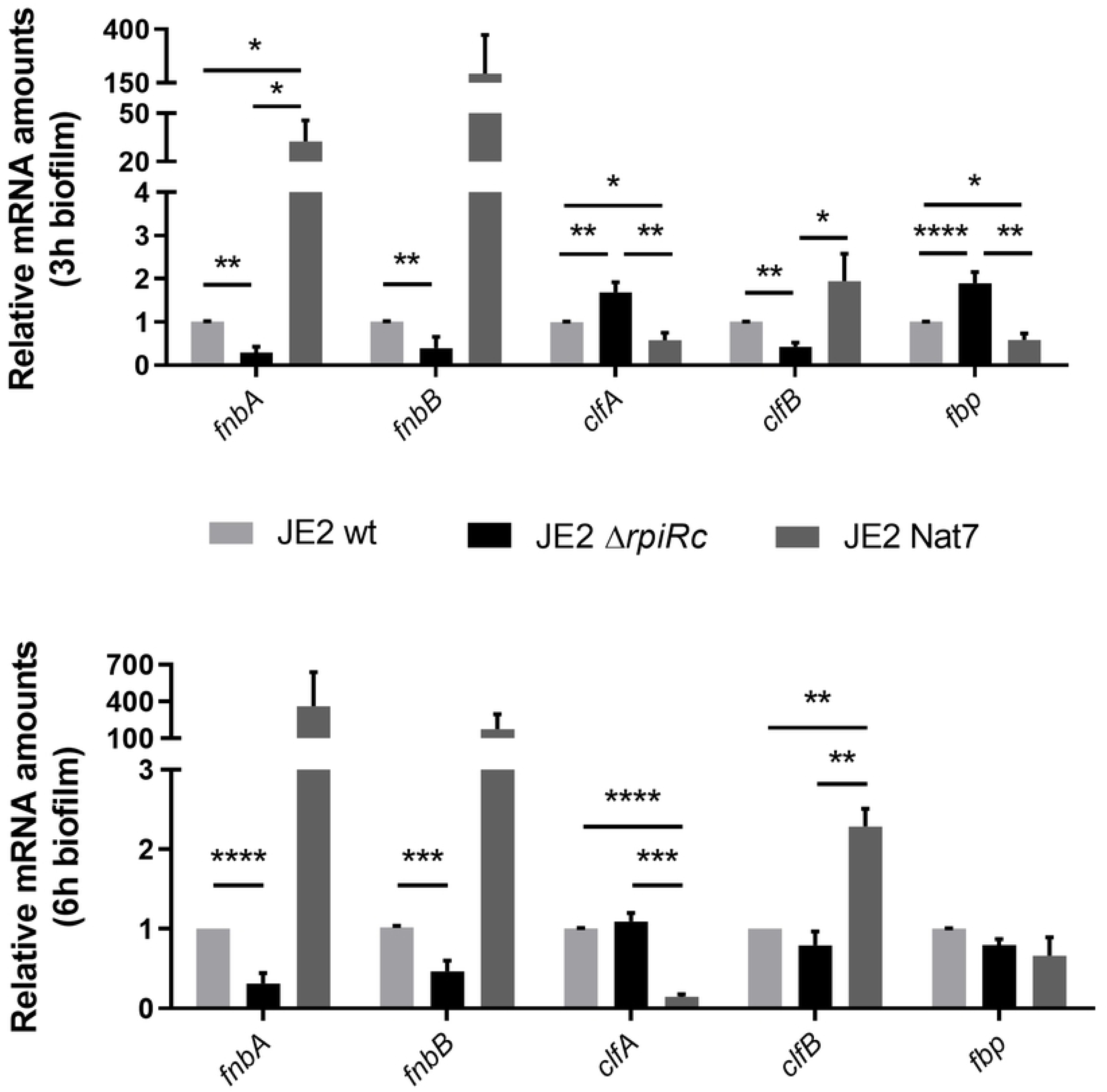
Relative MSCRAMMs mRNA expression in biofilms. (A) RpiRc dependent MSCRAMMs regulation in 3h old biofilms and (B) in 6h old biofilms. Gene expression was estimated by qRT-PCR from RNA, using 16S as gene expression normalization. For each gene, its expression in the wt strain was used as the reference mRNA amount (=1). Light grey bars represent USA300 JE2 wt strain, black bars represent USA300 JE2 Δ*rpiRc* strain and grey bars represent USA300 Δ*rpiRc* Nat7 strain. Data represents mean of six (A) or four (B) independent experiments; *; **; ***; ****: p<0.05; <0.01; <0.001; <0.0001 respectively.

To confirm that *fnbA-B* transcriptional regulation by RpiRc is directly linked to fibronectin binding protein production, USA300 JE2, UAMS-1, SA564 and SA113 wt and Δ*rpiRc* strains binding to fibronectin was tested (Figure 8). Binding to Fibronectin for strains UAMS-1, USA300 JE2 and SA564 was twice lower compared to SA113. *rpiRc* deletion in these three strains led to a two to three fold decreased adhesion to fibronectin. As observed and described previously in this manuscript for biofilm formation or eDNA amounts in the biofilm, *rpiRc* deletion does not alter SA113 binding to fibronectin suggesting that *rpiRc* dependent mature biofilm formation and MSCRAMMs transcriptional regulation are mediated by the same regulatory pathway, RsbU dependent. RpiRc regulates *fnbA-B* transcription leading to decreased FnBPA-B protein production.

**Figure 8:**
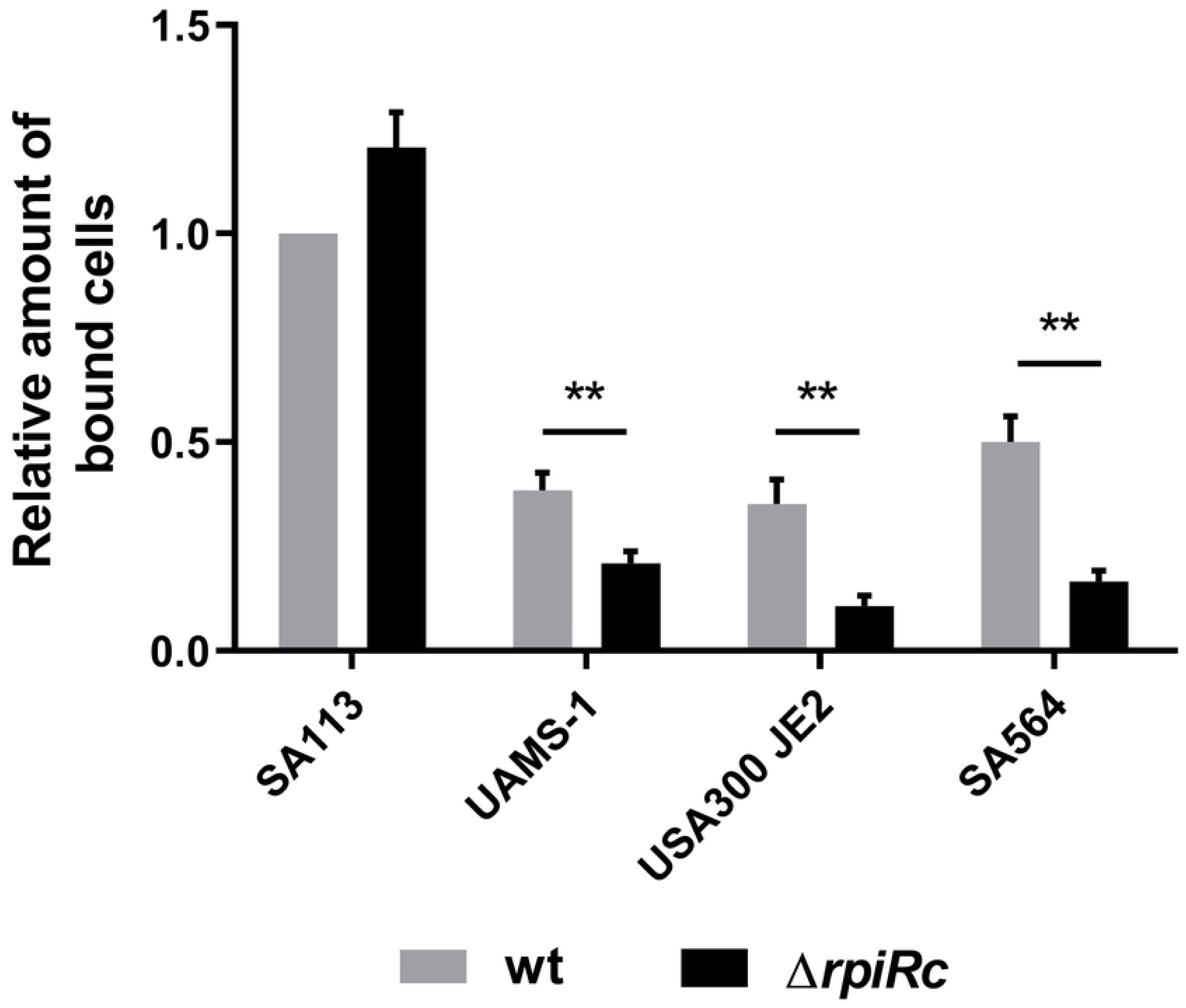
Adhesion to fibronectin requires RpiRc. UAMS-1, USA300 JE2, SA564 and SA113 wt and Δ*rpiRc* strains were incubated in wells precoated with fibronectin. The amount of bound bacteria to fibronectin was estimated using CV staining and normalized to SA113 wt (=1). Light grey bars represent wt strains and black bars represent Δ*rpiRc* strains. Data represents mean of six independent experiments; **: p<0.01.

### RpiRc enhance *ex vivo* and *in vivo* virulence

RpiRc regulates *fnbA-B* gene expression leading to increased FnbA-B protein production. Fibronectin binding proteins are key regulator of *S. aureus* ability to invade host cells. Transcription of ClfB which is involved in epithelial cell invasion is also under the control of RpiRc. Internalization experiment was performed in HEK293 cells with USA300 JE2, UAMS-1 and SA113 wt, Δ*rpiRc* and Nat7 strains (Figure 9). As positive and negative control Cowan (Figure 9A) and KH11 (Figure 9B) were used respectively (73, 74). Wild type strains USA300 JE2, UAMS-1 and SA113 interact and internalize inside HEK293 cells as much as the positive control Cowan, around 20% of HEK293 cells have internalized bacterial cells (Figure 9C). Internalization with UAMS-1 wt is lower compared to other wt strains. RpiRc deletion in UAMS-1 and USA300 JE2 presented decreased ability to interact and internalize inside HEK293 cells ranging from 5 to 8% of HEK293 cells respectively. UAMS-1 Δ*rpiRc* showed similar internalization into HEK293 cells as the negative control KH11, meaning that UAMS-1 Δ*rpiRc* is no longer able to internalize inside non-phagocytic cells (Figure 9C). Deletion of *rpiRc* in SA113 does modify internalization into HEK293 cells. Over-expression of *rpiRc* in USA300 JE2 and UAMS-1 restored internalization into HEK293 cells (Figure 9C). In contrary to *fnbA-B* expression which was highly increased in the complemented strains, internalization do not show *rpiRc* dose-dependent effect as complemented strains do not show increased internalization compared to wt or positive control. RpiRc regulation of fibronectin binding protein is directly related to *S. aureus* ability to interact and invade host cells.

**Figure 9:**
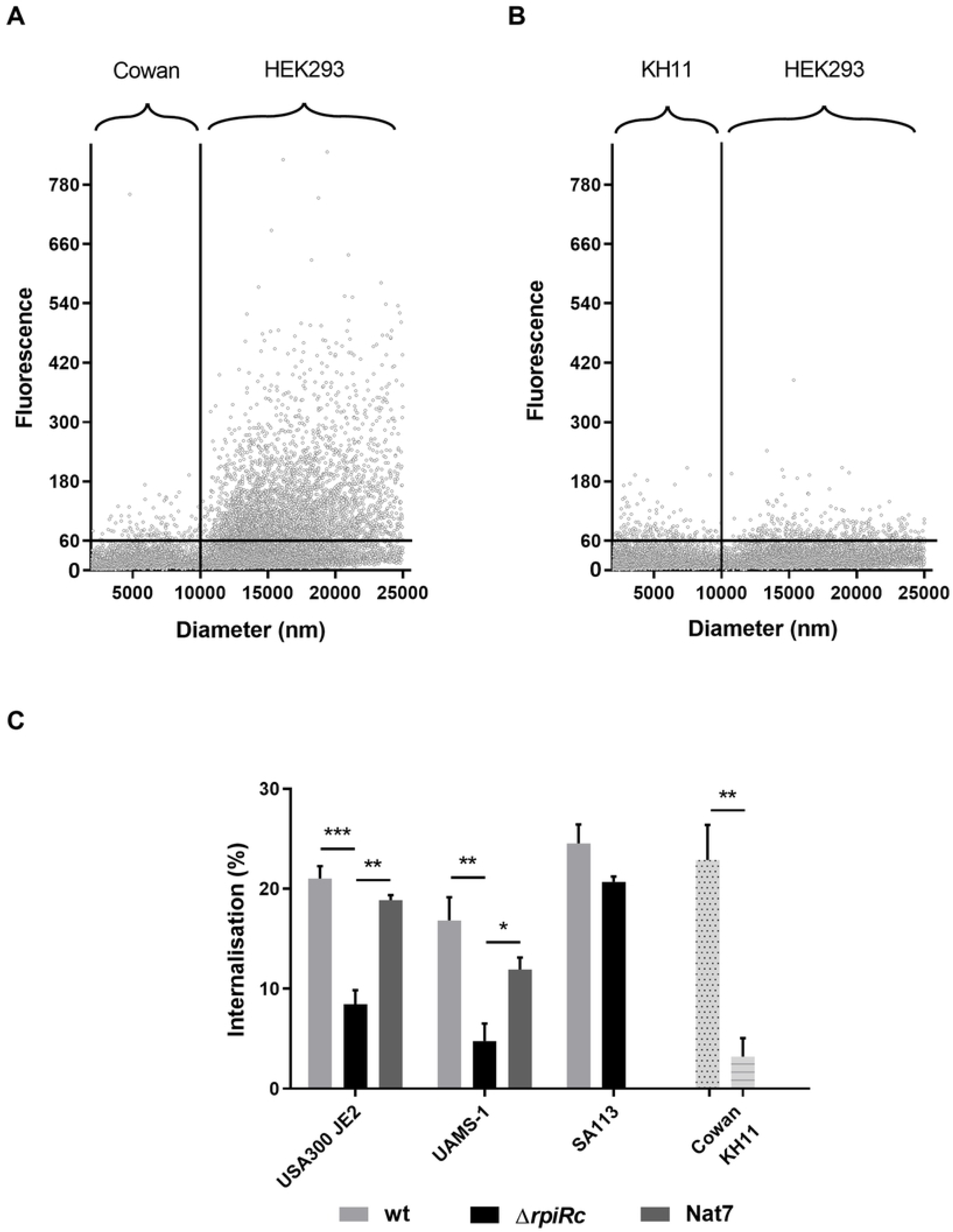
RpiRc is required for interaction and internalization into non-phagocytic cells HEK293. (A) Positive and (B) negative control for internalization using *S. aureus* Cowan and *S. epidermidis* KH11, respectively. Bacteria were considered fluorescent with a fluorescence higher than 60. HEK293 cells were considered bigger than 10µm (Figure S9). (C) Internalization into non-phagocytic cells HEK293 was performed with USA300 JE2, UAMS-1 and SA113 wt (light grey bars), corresponding Δ*rpiRc* (black bars) and complementation Nat7 (grey bars; except for SA113). Cowan (light grey bar with black dots) and KH11 (light grey bar with lines) were included as positive and negative controls for internalization, respectively. Invasiveness of *S. aureus* strains was determined as the percentage of fluorescent HEK293 cells (higher than 60 fluorescence units) among the whole population of HEK293 (size bigger than 10µm) as estimated in the MoxiFlow cytometer (A, B). Data represents mean of five (USA300 JE2 and UAMS-1 wt and Δ*rpiRc*), four (Cowan and KH11) or three (complemented strains and SA113 wt and Δ*rpiRc*) independent experiments; *; **; ***: p<0.05; <0.01; <0.001 respectively.

RpiRc plays a major role in the interaction with host cells *in vitro*. To test a role for RpiRc *in vivo*, USA300 JE2 wt and Δ*rpiRc* strains were inoculated in a subcutaneous catheter in mice. After 5 days, mice were sacrificed and the amount of biofilm in the catheter, the capsule and bacteria in the surrounding tissue were estimated as described in the material and method section. Biofilm formation in the catheter is very similar between wt and mutant strain for all the inoculum tested (Figure 10A). The amount of CFU recovered from the capsule showed more variation mostly for the mutant strain with an inoculum at 1.10^4^ (Figure 10B). No differences could be observed statistically significant between wt and mutant strain in the catheter and in the capsule. The amount of CFU/mL recovered for mutant strains in the capsule between inoculum 1.10^4^ and 2.10^5^ is not statistically different but present a trend for decreased CFU recovery with inoculum 2.10^5^. Further investigation would be required to confirm any increased capsule colonization of USA300 JE2 Δ*rpiRc* strain with an inoculum of 10^4^ CFU. USA300 JE2 wt strain requires a minimum inoculum of 10^4^ CFU to colonize host tissues (Figure 10C). The number of mice with this inoculum is low but the higher amount of mice with an inoculum of 6.10^4^ confirms the ability of USA300 wt to colonize tissues with a mean 1.10^3^ CFU/mg. The mutant strain is also able to colonize host tissues but requires a higher inoculum. No CFU counting is available with an inoculum of 10^3^. The mutant strain is not able to colonize host tissues with an inoculum of 10^4^ in contrary to wt strain (Figure 10C). Mutant strain requires an inoculum of 2.10^5^ CFU to colonize the tissue (significant difference between 10^4^ and 2.10^5^, p < 0.01). Tissue colonization with wt and *rpiRc* mutant strain is inoculum dependent (in contrary to catheter colonization), wt strains requiring an inoculum of 10^4^ CFU while Δ*rpiRc* requires an inoculum of 2.10^5^ CFU. RpiRc plays a major role in *S. aureus* virulence. It is required for adhesion and internalization in eukaryotic cell as well as for host tissue colonization in a mouse model of catheter infection.

**Figure 10:**
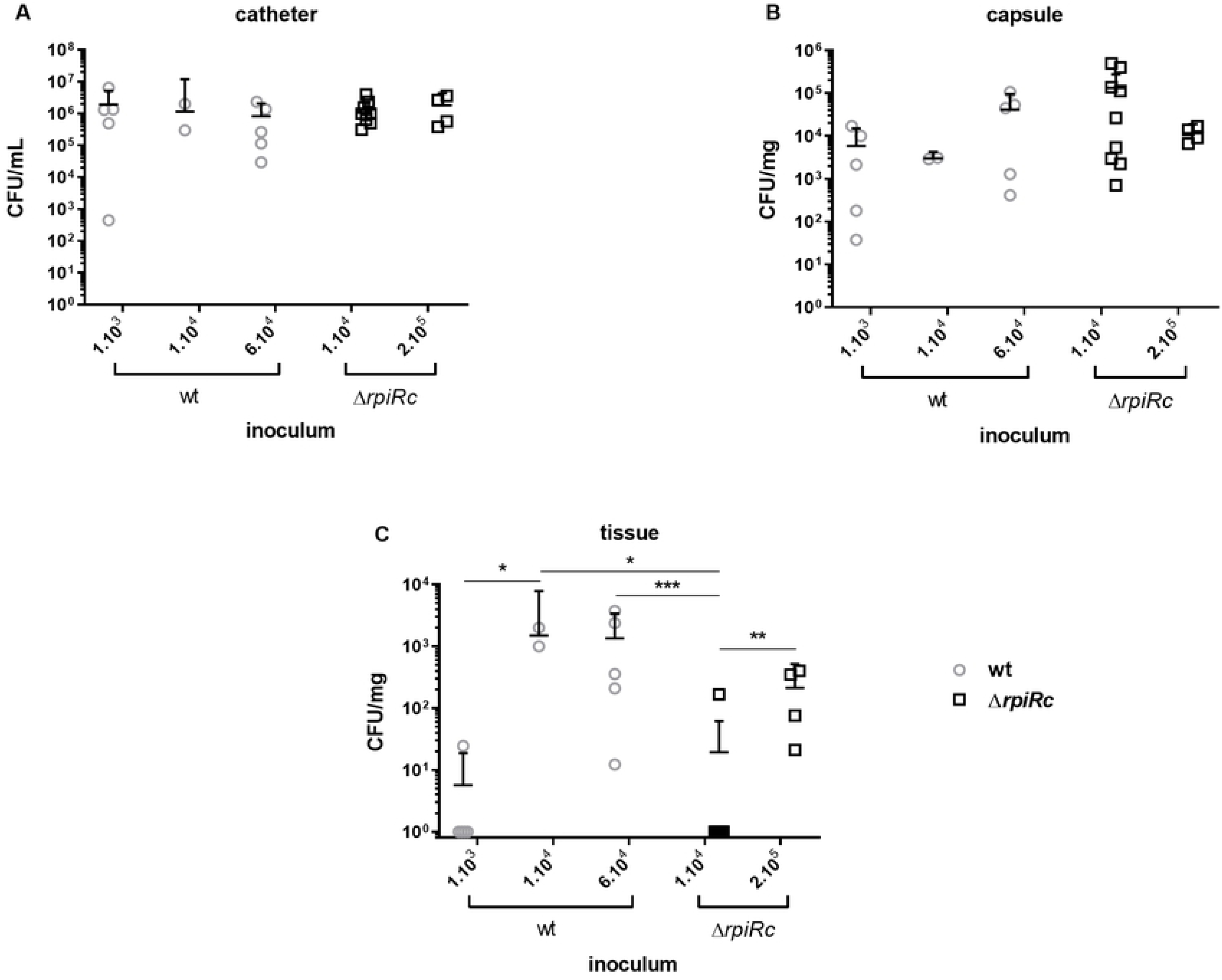
RpiRc is required for *in vivo* virulence in a mouse model of catheter infection. Different inoculum of USA300 JE2 wt and *rpiRc* mutant strains were introduced perioperatively in a catheter enclosed in a capsule. Catheter and capsule were explanted after 5 days and CFU counts were determined in the catheter (A), in the capsule (B) and in the surrounding tissue (C). Data represents means + 95% CI of 5, 2, 5, 9 and 4 independent experiments for wt with inoculum of 1.10^3^, 1.10^4^, 6.10^4^ CFU/catheter and Δ*rpiRc* with inoculum of 1.10^4^ and 2.10^5^ per catheter respectively; *; **; ***: p<0.05; <0.01; <0.001 respectively.

## Discussion

### RpiRc does not regulate PIA dependent biofilms

Using the recently available Nebraska Transposon Library, a new actor in the regulation of eDNA and biofilm formation of *S. aureus* could be identified. RpiRc is an enzyme involved in the pentose phosphate pathway (PPP), identified by homology from *E. coli* (75). It was first described in *S. aureus* as a regulator of RNAIII and virulence under planktonic conditions *in vitro* and *in vivo* respectively (75, 76). PPP starts from G6P to produce ribose required for nucleic acid synthesis. Without RpiRc, nucleic acid synthesis should decrease and growth should be affected. No difference in growth was monitored between wt and *rpiRc* mutant strains, suggesting that decreased eDNA might not related to decreased nucleic acid production in *rpiRc* deleted strains. *rpiRc* mutant strains secrete a compound in mature biofilms, under the control of RsbU, that degrades eDNA matrix from wt biofilms, confirming that decreased eDNA on *rpiRc* mutant biofilms is related to eDNA degradation rather than eDNA production. G6P is also required for glycolysis as well as for Glc-N-6-P production. This molecule is then metabolized into UDP-Glc-NAc which is required for peptidoglycan production but is also a precursor of the PIA (77). It was hypothesized that *rpiRc* could be involved in PIA dependent biofilm formation. RpiRc deletion in the PIA dependent biofilm producer SA113 as well as DispersinB addition to USA300 JE2 *rpiRc* mutant did not showed any modification in the biofilm formation compared to wt strain, strongly suggesting a PIA independent biofilm regulation by RpiRc.

It was reported that NCTC8325 strain and derivative have several mutations and should not be used as unique model for deletion experiments (78, 79). SA113 is a derivative strain from NCTC8325. It was observed in this laboratory strain SA113 a decreased planktonic growth when RpiRc is over-expressed in growing biofilm (6h). This decrease planktonic growth due to RpiRc overexpression is related to increase sessile/planktonic cell ratio. The decreased planktonic growth rate was also observed in the other strains but was less pronounced. RpiRc is an important regulator in bacterial cells and high amounts of this protein are probably toxic to the bacteria, strongly delaying planktonic growth, meaning that RpiRc could be targeted for antibacterial therapy. The side effect is an increased sessile/planktonic cells ratio.

### RpiRc involve RsbU dependent but *nuc* independent eDNA regulation for *S. aureus* biofilm formation

*S. aureus* virulence and biofilm formation is tightly regulated by SarA, Agr and SigB (56). They interact together and are all involved in the production or inhibition of extracellular proteins, regulating biofilm matrix stability. A previous publication showed that under planktonic growth conditions, RpiRc positively regulated all these factors (75). The laboratory strain SA113 contains a point mutation in *rsbU*, the regulator of SigB activity. The hypothesis was that absence of phenotype observed in SA113 Δ*rpiRc* is related to this mutated *rsbU*. RsbU deletion in other strains than SA113 confirmed results observed in this laboratory strain, RpiRc is involved in RsbU regulation to control biofilm formation and eDNA production/turn over in mature biofilms. An interaction between RpiRc and RsbU was previously suggested without phenotypic information (75, 76). From a review of the literature, this is the first report on RsbU directly involved in the regulation of eDNA amounts in mature *S. aureus* biofilms. It was shown previously by microarray experiment that *rsbU* deletion led to increased *nuc* expression and production but phenotype for eDNA amounts on biofilms was not experimented (80, 81), and we excluded *nuc* as an actor of eDNA regulation mediated by RpiRc.

RpiRc deletion is responsible for decreased eDNA and biofilm formation in mature biofilms by secreting an enzyme or a compound as suggested by the addition of the supernatant of USA300 JE2 Δ*rpiRc* mature biofilm to wt mature biofilm. The decreased eDNA recovery form the treated wt biofilms suggested active eDNA degradation, meaning over-production or stability of a nuclease. The major nuclease secreted and described in eDNA degradation in *S. aureus* biofilms is the thermonuclease Nuc1. Nuc2 is the second nuclease, similar to Nuc1 and also involved in eDNA degradation (to a lesser extend) to be described. This second nuclease is not secreted, bound to the membrane but active in the extracellular side of the bacteria. Both nucleases were deleted in SA564 Δ*rpiRc* strain without any modification of *rpiRc* mutant phenotype (neither for eDNA nor for biofilm formation). RpiRc regulates eDNA stability in a Nuc1-Nuc2 independent manner, meaning that another nuclease is involved in the degradation of eDNA in mature *S. aureus* biofilms under the control of RpiRc.

### Proteins involved in RpiRc dependent *S. aureus* biofilm formation

USA300 JE2 Δ*rpiRc* produces an extracellular compound degrading the extracellular matrix of the biofilm and more precisely eDNA. Analysis of the most concentrated protein fraction on an SDS acrylamide gel revealed increased production of many secreted protein and surface associated proteins involved in adhesion/biofilm formation or virulence. RsbU regulates SigB activity which is an inhibitor of extracellular proteases (82, 83). Two of these proteases, SspA and SspB, a serine and cysteine protease are co-transcribed and were increased in the supernatant of USA300 JE2 Δ*rpiRc*. SspA is able to degrade fibronectin binding proteins (84) and Spa (85). SspB is involved in increased virulence *in vivo* (83,86,87). The decreased eDNA amounts in *rpiRc* mutant biofilm is RsbU dependent, suggesting that increased protease production is not responsible for eDNA modification because it must be RsbU independent as previously demonstrated (82). Proteases are also positively regulated by Agr - RNAIII (82,87,88) and SarR (89). Proteases involvement in the biofilm formation of *rpiRc* mutant remains probable and would require additional investigations.

Other proteins involved in biofilm formation are the major autolysin Atl as well as MecA (67, 90). Atl regulates the amount of eDNA in the biofilm, like the hemolysin Hlb (91), also identified in the supernatant of USA300 JE2 Δ*rpiRc*. The absence of increased autolysis in JE2 Δ*rpiRc* suggests an Atl independent eDNA regulation. PrsA is a lipid anchored chaperone protein required for *S. aureus* resistance to vancomycin and oxacillin (92). The amount of PrsA in the supernatant of JE2 Δ*rpiRc* is lower than JE2 wt supernatant suggesting decreased resistance to glycopeptide and oxacillin of *rpiRc* mutant strains, but was not confirmed by MIC experiment. The second immunoglobulin-binding protein (Sbi), is found anchored in the membrane and extracellularly (93). Sbi is a pro-inflammatory staphylococcal antigen (94) and both forms contribute to *S. aureus* immune evasion (phagocytosis and Neutrophil killing) (93). In addition to the increased protease and hemolysins production observed, decreased production of Sbi in JE2 Δ*rpiRc* would suggest increased virulence for this mutant *in vivo* as previously observed in two *in vivo* mouse models of abscess and pneumonia using planktonic bacteria (75, 76).

eDNA regulation by RpiRc involves nucleic acid degradation by a secreted compound altering mature biofilm stability. Neither Nuc1 nor Nuc2 are involved in this regulatory pathway. Other nucleases were identified in the supernatant of mature *rpiRc* mutant biofilms. AdsA weight is too high for the band analyzed. The second nuclease identified is Sln and was selected for further mutagenesis experiment. Sln deletion in USA300 JE2 Δ*rpiRc* suggested that RpiRc regulates eDNA amounts in mature biofilms by inhibiting this protein, but only on growing biofilms and not mature biofilms. To confirm a role of Sln in the regulation of eDNA by RpiRc, additional investigation would be required. Sln is a 5’-nucleotidase described in *Haemophilus influenzae* degrading extracellular riboside and involved in the utilization of nicotinamide mononucleotide (95, 96) but also in the degradation of nucleotides in *Vibrio parahaemolyticus* to cleave extracellular ATP and use the resulting adenoside (97, 98). The consequence of this intracellular nucleotide regulation is a decreased eDNA amounts in the matrix of mature biofilms resulting in decreased biofilm stability, meaning decreased amount of mature biofilm compared to the parental strain.

Previous work showed that RpiRc positively regulated the production of RsbU in planktonic cells. The present work demonstrated that RpiRc regulates RsbU transcription and production under biofilm formation conditions leading to increased eDNA stability on the matrix, meaning increased eDNA amounts correlating with increased recalcitrance to different classes of antibiotics. RsbU deletion also restored SA564 wt biofilm amounts in *rpiRc* deletion strain, but did not reached the highest amount as observed in the single *rsbU* deletion strain. In UAMS-1, *rsbU* deletion showed decreased biofilm formation while double mutant produced more biofilm than single *rpiRc* mutation; it did not reached wt amounts, suggesting that other factors must be involved in RpiRc dependent biofilm regulation.

### Regulation of MSCRAMMs

Decreased biofilm formation observed in *rpiRc* mutant strains is not consistent with increased RsbU production as RsbU is an inhibitor of *agr* and *RNAIII* transcription, both related to decreased biofilm formation. Gaupp and colleagues showed increased *RNAIII* / *agr* expression in planktonic *rpiRc* mutant strains. The expression level of these genes under biofilm conditions, and assessed by qRT-PCR, presented increased amount of *agr*, *RNAIII* and *hla* mRNA as observed in planktonic conditions (76). The QS and RNAIII are involved in the inhibition of biofilm formation, consistent with decreased biofilm formation in *rpiRc* mutant strains. This increased expression is visible after 5h of growth and the highest effect of biofilm decrease was observed on mature biofilms, meaning that another factor may be involved in the regulation of biofilm formation through this pathway. Indeed, Agr and RsbU control the expression and production of membrane associated proteins, the MSCRAMMs.

### RpiRc regulates FnbA-B transcription and production

MSCRAMMs are composed by many surface associated proteins like the fibronectin binding proteins A and B (FnBPA-B) or the clumping factors A and B (ClfA-B) (99, 100). It was recently reported that FnBPA domains involved in Fibronectin binding and biofilm formation are distinct and exclusive (38,101,102). Deletion of Fibronectin binding proteins did not alter cell adhesion and early biofilm formation on polystyrene surfaces, but altered intercellular interactions (102). RpiRc dependent *RNAIII* increased transcription only led to decreased *fnbA-B* transcription under biofilm formation conditions, explaining the absence of early biofilm formation modification on polystyrene surfaces. Decreased intercellular interactions related to decreased *fnbAB* transcription could be a second mechanism under the control of RpiRc to regulate biofilm formation, and is consistent with decreased mature biofilm formation. We speculate that eDNA might bind to FnBPA-B, decreased FnBPA-B production leading to decreased eDNA assembly on bacterial surface and decreased biofilm formation.

*RNAIII* inhibits the expression of the fibronectin binding protein A and B (103) and Agr inhibits *clfB* expression through rot regulation (64). These proteins are required for fibronectin and fibrinogen binding as well as for biofilm formation (38, 104). *rpiRc* deletion was responsible for decreased *fnbA-B* and *clfB* transcription. Consistent with previous observations (105), decreased *fnbA-B* transcription led to decreased binding to fibronectin confirming that FnBPA-B production is affected in the *rpiRc* mutant. However, SA113 Δ*rpiRc* was not affected in fibronectin binding, confirming that RpiRc dependent *fnbA-B* and *clfB* transcription is also under the control of RsbU-SigB. SigB is an inhibitor of RNAIII transcription (106), which does not correlate with *RNAIII* and *fnbA-B* - *clfB* transcription levels observed in the biofilms of *rpiRc* mutants. SigB also enhances *fnbA* transcription in strain Newman (80). While RsbU-SigB is an inhibitor of the *agr* system (106), SigB activates *sarA* transcription (59). SarA production and transcription are enhanced in the *rpiRc* mutant under planktonic conditions (75). SarA is a positive regulator of *fnbA-B* transcription (107) but also a positive regulator of the *agr*/RNAIII transcription (108), meaning that RpiRc regulates *fnbA-B* transcription through the inhibition of RsbU leading to decreased *sarA* transcription, decreased *agr* transcription, decreased RNAIII transcription leading to decreased *fnbA-B* inhibition, increased FnBPA-B production and increased binding to fibronectin and biofilm formation.

### RpiRc is required for *ex vivo* cell adhesion

FnBPA-B are not only important for biofilm formation but also for bacterial interaction with host cells. While FnBPA-B are not involved in biofilm formation on catheter (109), non-phagocytic host cells invasion by *S. aureus* requires fully functional FnBPs (110–112), FnBPA-B being sufficient for internalization (110). FnBPA cooperate with ClfA for *in vivo* invasion and colonization. ClfA is only required for colonization, invasion in absence of FnBPA being impossible (112). Similarly, ClfB is required for epithelial cell invasion and *clfB* expression decreased in the *rpiRc* mutant strains. The ability of *rpiRc* mutants to interact and invade HEK293 cells was tested. RpiRc is required to invade host cells and RpiRc deletion almost abolishes it. Once again, *rpiRc* deletion in SA113 did not alter SA113 competence to invade HEK293 cells, confirming that *S. aureus* ability to invade host cells is under the control of RsbU. *In vitro*, *rpiRc* deletion causes decreased biofilm formation related to decrease eDNA amounts. *In vivo*, similar amounts of biofilms were produced in the catheter compared to USA300 JE2 wt strain and a trend for increased biofilm formation of USA300 JE2 Δ*rpiRc* in the capsule was noticed, due most probably to increased *clfA* expression in the biofilm of USA300 JE2 Δ*rpiRc* strain.

### *In vivo* biofilm and virulence model

*In vivo* biofilm formation using a mouse model of catheter infection did not reveal differences between wt and *rpiRc* mutants, in contrary to *in vitro* biofilm formation. Gaupp and colleagues demonstrated that *rpiRc* deletion resulted in increased virulence using a mouse model of abscess and pneumonia (75). In the mouse model of catheter infection, difference could only be observed in the tissues, where USA300 JE2 Δ*rpiRc* presented decreased colonization, in contrary to previous *in vivo* observation (75) and to *in vitro* biofilm and planktonic increased expression of *agr* and *RNAIII* ((113) and present study). In this study, RpiRc was shown to be required for adhesion and internalization in non-professional phagocytes cells HEK293, suggesting decreased virulence in the mutant strains in an RsbU dependent manner. RsbU is required for *S. aureus* virulence for induced arthritis and sepsis in mice (114) but not in a murine subcutaneous skin abscess model of infection (82), while decreased *fnbA-B* is related to decreased *in vivo* virulence (115). Increased virulence of *rpiRc* mutants observed by Gaupp and colleagues (75) and decreased virulence observed in the catheter suggests two different roles for RpiRc dependent on the mode of growth of the bacteria, e.g. decreased virulence when grown in a biofilm. It is most probable that increased Agr and *RNAIII* under biofilm mode of growth only regulates surface proteins, leading to *in vitro* decreased protein dependent biofilm formation and *ex vivo* – *in vivo* decreased virulence without affecting biofilm formation *in vivo*. Decreased ability of USA300 JE2 Δ*rpiRc* to invade host tissue in the mouse model of catheter infection could be related to decreased expression of *fnbA-B* in this mutant strain. On the other hand, increased *clfA* expression in the mutant strain supports the similar colonization on the catheter and even the increased colonization on the capsule compared to the wt strain.

## Conclusion

RpiRc is an important regulator of biofilm formation in *S. aureus*. In mature biofilms, RpiRc regulates in an RsbU dependent manner protein or metabolite secretion involved in eDNA stability. One target protein is Sln, allowing the bacteria to digest eDNA and use riboside to overcome the potential lack of nucleotide synthesis. RpiRc also regulates the expression and production of fibronectin binding proteins and clumping factor B, leading to increased adhesion to fibronectin, increased cell interaction under biofilm formation conditions and increased internalization and invasion with host cells (Figure 11). Regulation of eDNA on mature biofilm and regulation of FnBPA-B / ClfB depict two mechanisms under the control of RpiRc to regulate *in vitro* and *in vivo* biofilm formation in *S. aureus*, increasing its virulence.

**Figure 11:**
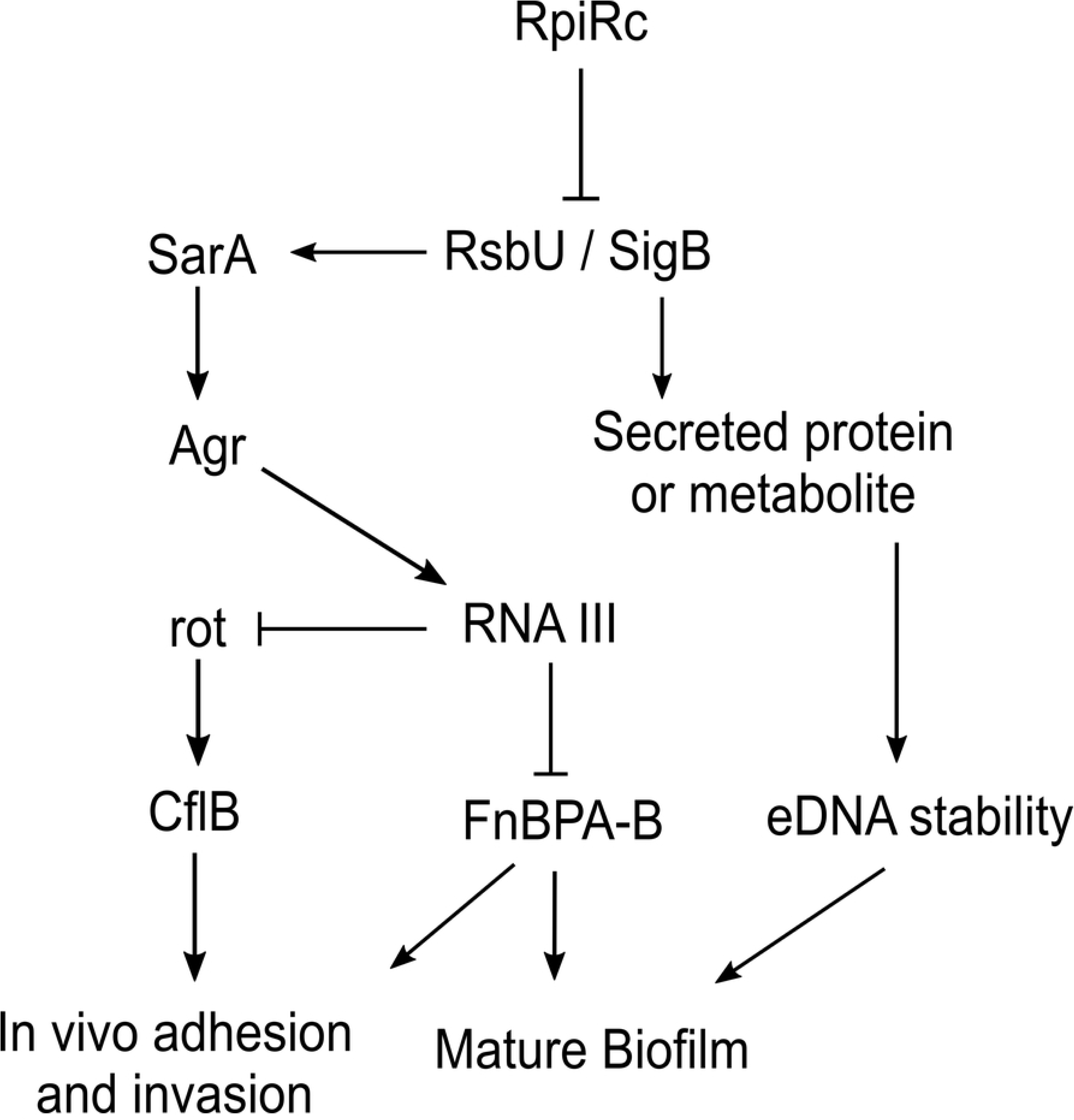
RpiRc regulatory pathways. RpiRc inhibits RsbU transcription and production leading to decreased free activated SigB. Inhibition of SigB result in decreased *sarA* transcription (59) decreased *agr* transcription and RNAIII inhibition (106). RNAIII repression leads to decreased *fnbAB* and *clfB* inhibition (64, 103) favoring biofilm formation and host cells invasion. On the other side, the secreted protein or metabolite under the positive control of RsbU involved in eDNA stability is inhibited, increasing eDNA amounts in biofilm matrix and increasing biofilm stability.

## Materials and methods

### Strains and conditions

Strains used in this study are presented in table 3. All wild type strains were grown on Mueller-Hinton (MH) agar plates at 37°C. *S. aureus* liquid cultures were in TSB supplemented with 1% glucose (TSBg) at 37°C with shaking (180rpm). *E. coli* DH5α was used for plasmid propagation, *S. aureus* RN4220, USA300 JE2, UAMS-1, SA113 and SA564 for deletion and complementation assays and *S. aureus* Cowan and *S. epidermidis* KH11 as control for internalization assay.

**Table 3:**
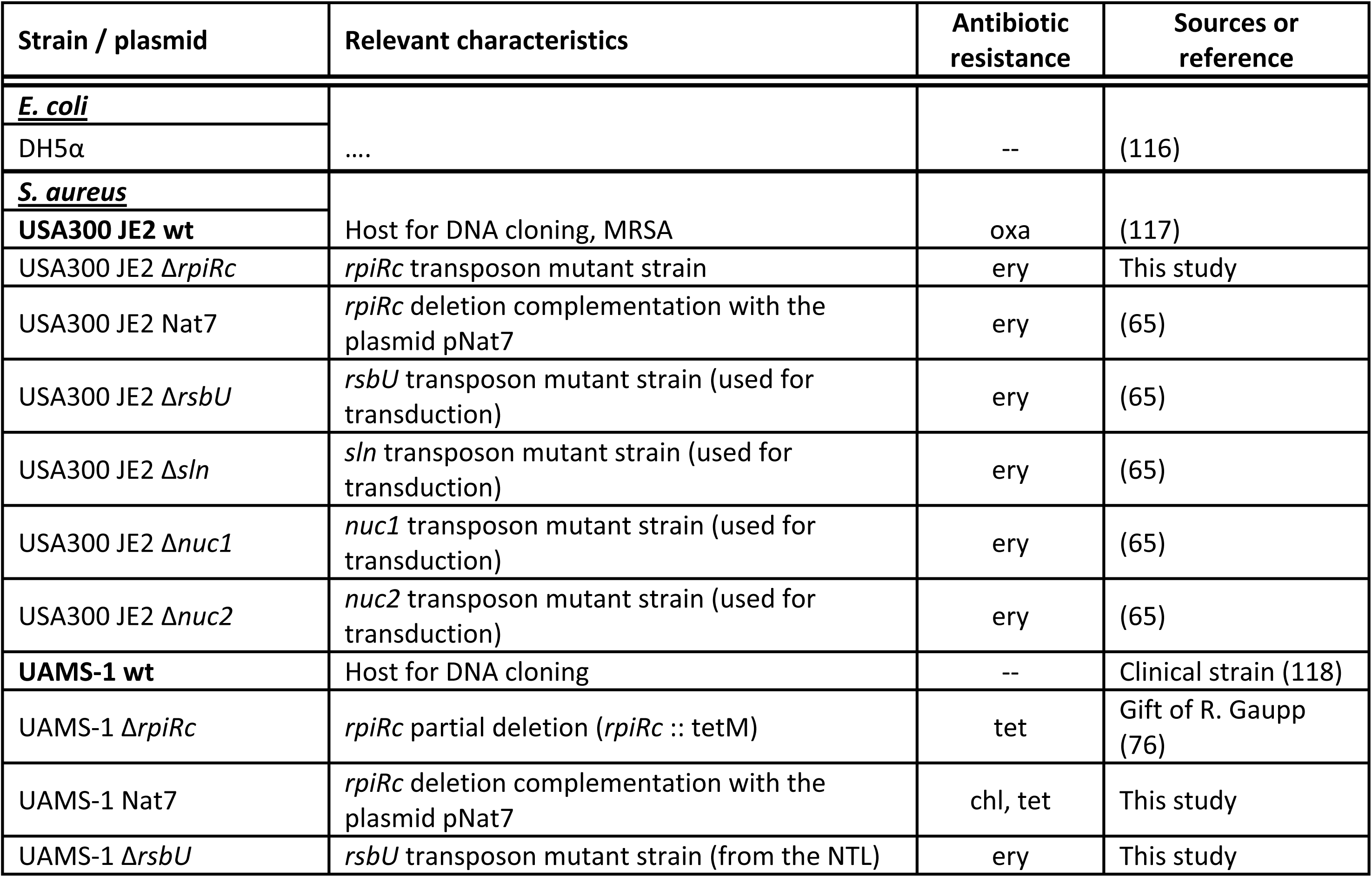
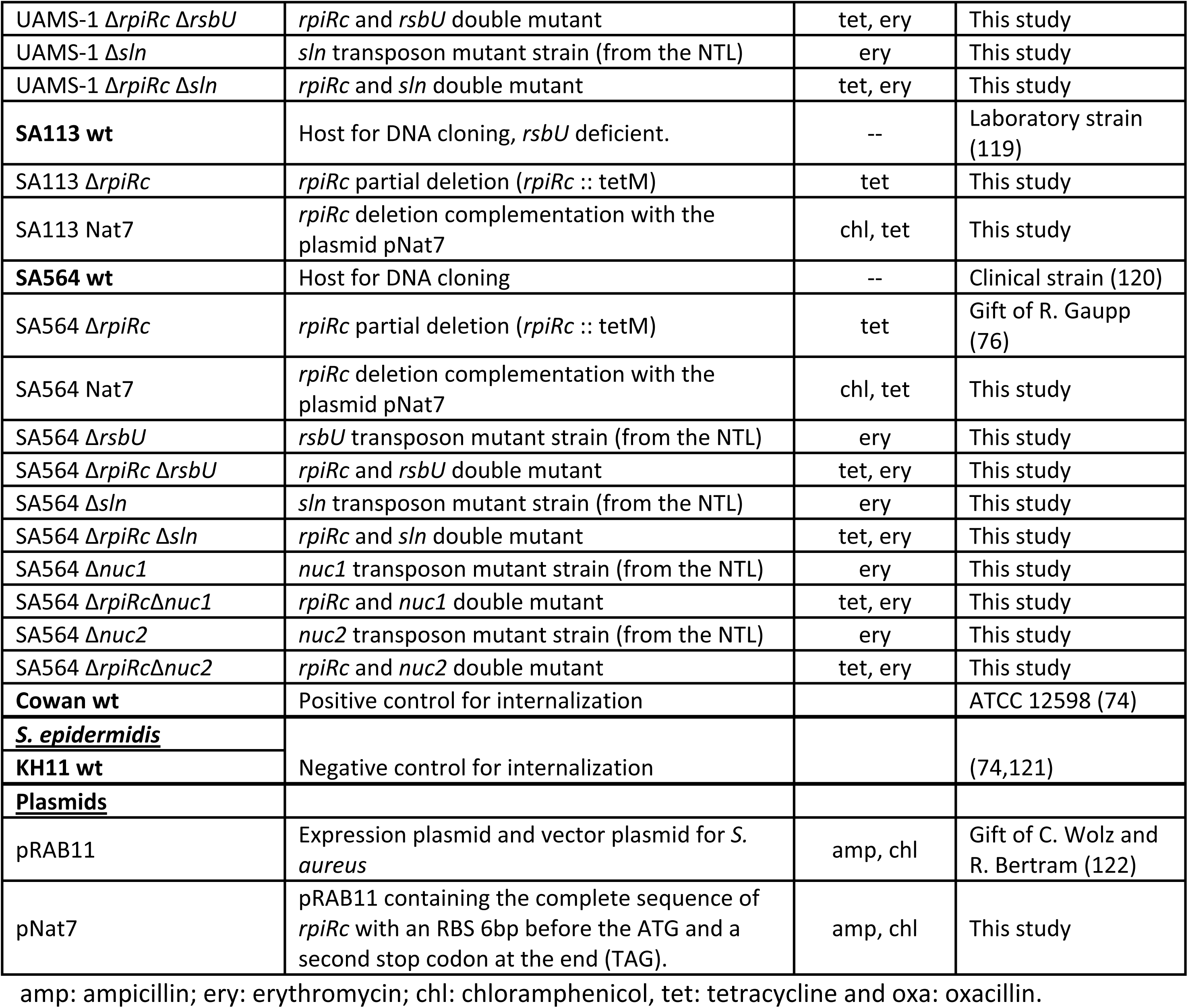
Bacterial strains and plasmids used in this study.

*S. aureus* mutant strains, and those containing plasmids, were plated on MH agar or grown in TSBg with 10µg/mL erythromycin, 10µg/mL tetracycline or 10µg/mL chloramphenicol (Table 3). For the complementation strains Nat7, 250ng/mL anhydrotetracycline were added to induce *rpiRc* expression and production.

### Mutant and complementation construction

*rpiRc* deletion in USA300 JE2 was obtained from the NTL and back-transduced using phi11 (54). UAMS-1 Δ*rpiRc* was kindly provided by Dr Gaupp (76) and was used as donor to delete *rpiRc* in SA564 and SA113 using phi11 transduction. RpiRc deletion was also back-transduced in UAMS-1. All other single (*rsbU*, *nuc1*, *nuc2*, *sln*) and double mutants (with *rpiRc*) were obtained by transduction from NTL mutants in UAMS-1 and SA564 backgrounds. Each mutant obtained in the lab (except *rpiRc*) was selected twice and tested together during the first replicate. Once the first experiment showed similar result between the two identical mutants, only one was kept for the rest of the experimental procedures.

RpiRc complementation was obtained using the pRab11 plasmid containing an anhydrotetracyclin inducible promoter. The complete sequence of *rpiRc* was amplified with modified forward primers (table 4; rpiRc_HpaI-RBS_F: 5’-ACCAAA*<u>GTTAAC</u>*AGGAGGTAAAAAA**<u>ATG</u>**ATTAATATGTCAAACGTA-3’) containing an RBS just before the ATG and a reverse primer with additional stop codons (table 4; RpiRc_EcoRI_R: 5’-TGATA*<u>GAATTC</u>***<u>CTATTA</u>**ATGTTTAAAGTTAATATTTGATAAATGTTTACGATAGTTAT-3’). This plasmid was introduced in USA300 JE2 Δ*rpiRc*, UAMS-1 Δ*rpiRc*, SA564 Δ*rpiRc* and SA113 Δ*rpiRc* using previously published protocol (54). Briefly, the ligated plasmid (Figure S10) was introduced in *E. coli* DH5α by heat shock. After purification, it was electroporated in *S. aureus* RN4220 and transduced with phi11 in the different target strains (Table 3).

**Table 4:**
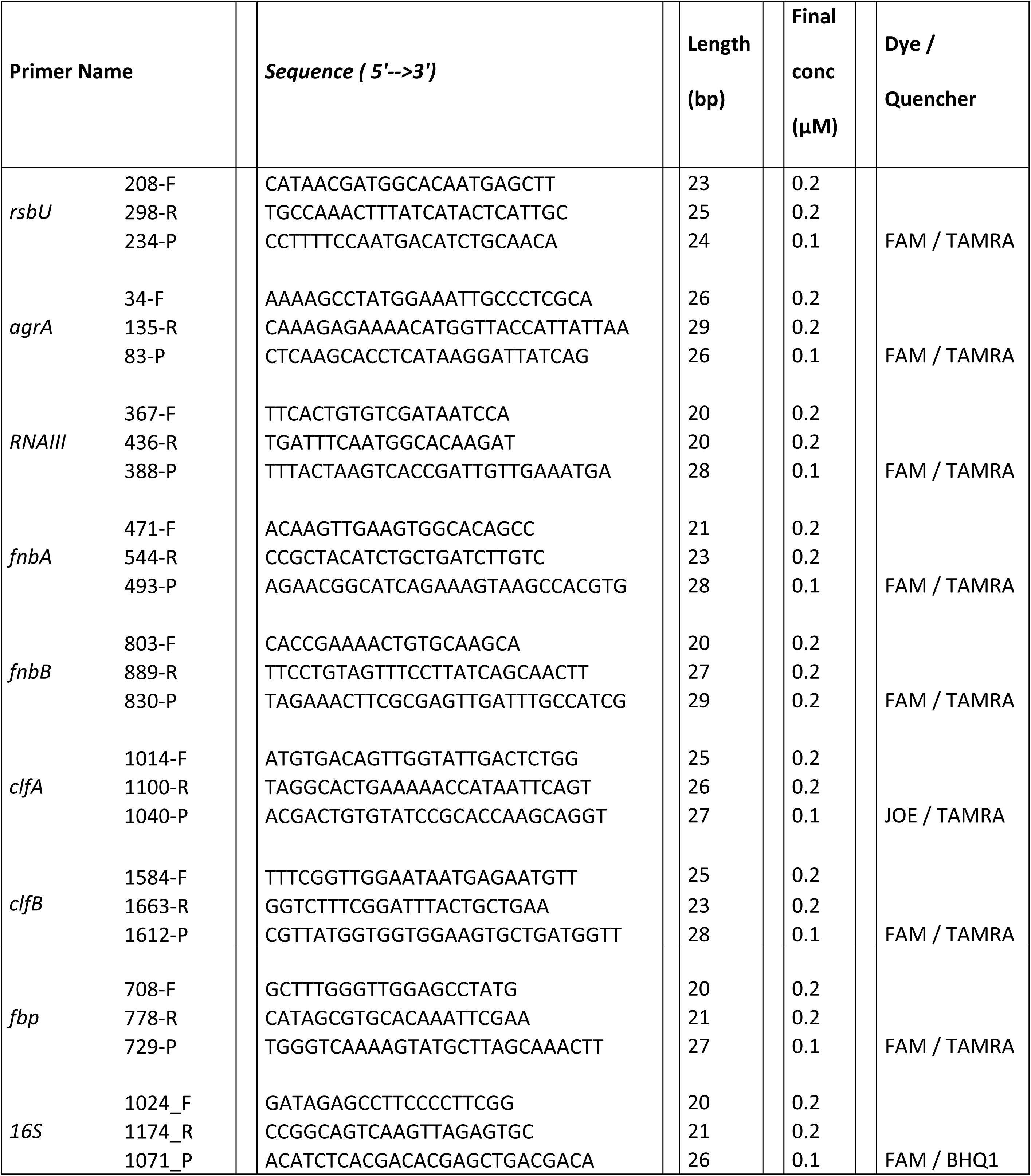
Primers and probes used in qRT-PCR.

### NTL screening

All the mutant strains from the mutant library were first re-isolated on MH + 10µg/mL erythromycin and frozen at −80°C individually. Due to the high number of strains to test, overnight cultures were performed in TSB + 1% glucose (TSBg) in 96 well plates at 37°C with slow agitation (100rpm). Dilutions 1/100 were performed similarly for all strains in TSBg in a 96 well plate and incubated 24h at 37°C without agitation. Biofilm formation was estimated using previously published protocol (54) using crystal violet staining. Biofilms were normalized using the median value of each column and row. This normalization method was validated using a plate with only wt strain USA300 JE2.

eDNA and PIA production in the biofilms of selected mutants were also tested in 96 well plates, following similar protocol as described above with addition of DNaseI and DispersinB at the beginning of biofilm formation (after dilution 1/100) as previously described (54). Modification of biofilm formation in presence of DNaseI correlated with the amount of eDNA present in the biofilms (54).

### Biofilm formation

All biofilm formation experiments after NTL screening were performed in 24 well plates at different time points (1.5h, 6h and 24h), following previously described protocol (54), without additional normalization.

### eDNA quantification

eDNA amounts were quantified from preformed biofilms at different time points (1.5h, 6h and 24h) using previously published protocol (54).

### eDNA addition to the biofilms

eDNA originating from SA113 Δ*gdpS* was purified (above protocol) (54) and added to culture medium (TSBg) at the beginning of USA300 wt and Δ*rpiRc* biofilm formation in 24 well plates. Plates were incubated 1.5h, 3h, 6h and 24h at 37°C without agitation. eDNA was purified from biofilms and quantified using a specific primer pair for the erythromycin cassette contained in the exogenous eDNA (SA113 Δ*gpdS*; Table 4) that is not present in USA300 JE2 wt or Δ*rpiRc*. The absence of non-specific amplification (USA300 JE2 Δ*rpiRc* carry a different erythromycin cassette) was tested using eDNA extracted from both wt and *rpiRc* mutant strain in absence of exogenous eDNA and showed no amplification. Similarly, exogenous eDNA was quantified from the SN to evaluate the part of this eDNA not bound or unbound to the biofilm.

### Supernatant (SN) collection and testing on biofilms

Supernatant of 24h old USA300 JE2 wt and Δ*rpiRc* biofilms were collected, centrifuged 10 minutes at 6000g and passed through a 0.22µm filter to eliminate any residual bacteria. The SNs collected were directly added to preformed USA300 JE2 mature biofilms (24h) after removal of its supernatant. As a control, one well with USA300 JE2 wt mature biofilm was not modified (no SN removal or replacement). Biofilms were then re-incubated at 37°C without agitation for additional 15 minutes, 1h, 6h and 24h and eDNA quantified.

### 24h biofilm SN protein anaylsis

#### Protein collection

Supernatant from 6 hours old USA300 JE2 wt and Δ*rpiRc* biofilms were collected and filtered with a syringe with a 0.22µm filter. Protease activity was inhibited using protease inhibitor cocktail tablets (cOmplete Tablets Mini; Roche Diagnostics, Indianapolis, USA). SN were then incubated 30min at 4°C in a solution of 20% (w/v) Trichloroacetic acid (TCA), followed by 1 hour centrifugation (13000g, 4°C). Pellets were then washed three times with ice cold acetone and resuspended in a 2x Laemmli solution before SDS page gel run.

#### In Gel Digestion

Band of interest were cut out from a 1D gel electrophoresis for digestion with trypsin by using the following procedure. Gel pieces were first destained by incubation in 200μL of 50mM ammonium bicarbonate and 30% acetonitrile (CH3CN) for 30 min at room temperature. Destaining solution was removed and gel pieces were then incubated for 35 min. at 56°C in 150μL of 10mM DTT in 50mM ammonium bicarbonate. DTT solution was then replaced by 150μL of 55mM iodoacetamide in 50mM ammonium bicarbonate and the gel pieces were incubated for 30 min at room temperature in the dark. Gel pieces were then washed for 30 min with 150μL of 50mM ammonium bicarbonate and for 30 min with 150μL of 50mM ammonium bicarbonate and 30% CH3CN. Gel pieces were then dried for 45 min in a Centrivap vacuum centrifuge (Labconco, Kansas City, USA). Dried pieces of gel were rehydrated for 45 min at 4°C in 50μL of a solution of 50mM ammonium bicarbonate containing trypsin at 6.25ng/μL. Extraction of the peptides was performed with 70μL of 1% TFA for 30 min at room temperature with occasional shaking. The TFA solution containing the proteins was transferred to a polypropylene tube. A second extraction of the peptides was performed with 70μL of 0.1% TFA in 50% CH3CN for 30 min at room temperature with occasional shaking. The second TFA solution was pooled with the first one. The pooled extracts were completely dried by evaporation under vacuum.

#### MS analysis

Samples were diluted in 10μL of loading buffer (5% CH3CN, 0.1% FA) and 4μL was injected on column. LC-ESI-MS/MS was performed on a Q-Exactive Hybrid Quadrupole-Orbitrap Mass Spectrometer (Thermo Fisher Scientific) equipped with an Easy nLC 1000 system (Thermo Fisher Scientific). Peptides were trapped on a Acclaim pepmap100, C18, 3μm, 75μm × 20mm nano trap-column (Thermo Fisher Scientific) and separated on a 75μm × 500mm, C18, 2μm Easy-Spray column (Thermo Fisher Scientific). The analytical separation was run for 90 min using a gradient of H2O/FA 99.9%/0.1% (solvent A) and CH3CN/FA 99.9%/0.1% (solvent B). The gradient was run as follows: 0-5 min 95 % A and 5 % B, then to 65 % A and 35 % B for 60 min, and 10 % A and 90 % B for 25 min at a flow rate of 250nL/min. For MS survey scans, the resolution was set to 70000 and the ion population was set to 3 × 106 with an m/z window from 400 to 2000. For protein identification, up to fifteen precursor ions were isolated and fragmented by higher-energy collisional dissociation HCD. For MS/MS detection, the resolution was set to 17000, the ion population was set to 1 × 105 with an isolation width of 1.6 m/z units. The normalized collision energies were set to 27%.

#### Protein identification

Peak lists (MGF file format) were generated from raw data using the MS Convert conversion tool from ProteoWizard. The peaklist files were searched against the StaphUSA300 database (uniprot sptr, release-2016-09, 5954 entries) using Mascot (Matrix Science, London, UK; version 2.5.1). Trypsin was selected as the enzyme, with one potential missed cleavage. Mascot was searched with a fragment ion mass tolerance of 0.020 Da and a parent ion tolerance of 10.0 ppm. Variable amino acid modifications were oxidized methionine. Fixed amino acid modification was carbamidomethyl cysteine. The mascot search was validated using Scaffold 4.5.1 (Proteome Software). Protein identifications were accepted if they could be established at greater than 95.0 % probability and contained at least 2 identified peptides.

### RNA extraction from 3 and 6-hour-old biofilms

For RNA extraction USA300 JE2 wt, Δ*rpiRc* and Nat7 were grown under biofilm condition as previously described (54) in 6 well plates for 3 and 6 hours. Then biofilms were washed with PBS once (3h) or twice (6h) and fixed with a solution of acetone ethanol (1/1, v/v). Then, bacterial biofilms were scrapped from the wells and bacterial RNA was extracted using RNeasy plus mini Kit (Qiagen Inc., Valencia, CA) following manufacturer’s recommendations. RNA purity and yield were evaluated with the Bioanalyzer (Agilent) and Nanodrop (Witec ag). The absence of DNA was verified by performing qPCR without RT.

### Real-Time Polymerase Chain Reaction (qRT-PCR)

MSCRAMMs mRNAs present in USA300 JE wt, Δ*rpiRc* and Nat7 were quantified. Primers and probes for 16S were used to standardize comparisons, as previously described (75, 76).

All experiments were performed with a one-step enzymatic kit in a Stratagene Mx3005P qPCR system (Agilent Technologies). For this experiment Brilliant III ultra-fast qRT-PCR master mix (Agilent) was used with ROX following manufacturer’s recommendations. Each probe was assessed for linearity by testing serial RNA dilutions. Primers and probes used in this experiment are listed in table 4.

### Fibronectin binding assay

Adhesion of bacterial suspensions to solid phase human fibronectin was assessed in a microtiter plate assay following recently published protocol (123) using TSBg instead of MHB.

### Internalization

*S. aureus* internalization in non-phagocytic cells HEK293 (adenovirus type 5 DNA-transformed primary human embryonic kidney, obtained from ATCC CRL-1573) was tested as previously described (73) with the following modifications. Living bacterial strains USA300 JE2 and UAMS-1, wt, Δ*rpiRc* and Nat7, SA113 wt and Δ*rpiRc*, Cowan and KH11 were stained with TRITC following previously published protocol (74). 1.10^6^ Stained bacteria were incubated 2 hours in presence of HEK293 cells (wells were coated with poly-ornithine prior to HEK293 addition) in 6 well plates. Then HEK293 cells were trypsinized, and internalization was estimated using a MoxiFlow cytometer (Witec AG, Switzerland). The smallest size (10µm to be considered as HEK293 cells was determined using the MoxiFlow cytometer with HEK293 cells alone (Figure S9). The lowest fluorescence intensity (fluorescence units; FU) to be considered as HEK293 containing a fluorescent bacteria (60 FU) was determined using the negative control for internalization KH11 fluorescent bacterial cells.

### Mouse model of catheter infection

Female C57BL/6 mice were kept in the animal facility of the Department of Biomedicine, University Hospital Basel. Animal guidelines were followed in accordance with the regulations of Swiss veterinary law (permit no. 1957). Under anesthesia, a 3-4 mm incision was made lateral to the spine, and a 1 cm catheter segment (Venflon™ Pro Safety, 14GA, Becton Dickinson) was inserted subcutaneously. Before closing the incision with a wound clip, 25 µL of pyrogene-free saline containing different inoculum of MRSA JE2 wt or Δ*rpiRc* was injected into the catheter. After 5 days, mice were sacrificed and catheter, surrounding capsule and tissue were aseptically removed. Catheter were transferred into eDNA extraction buffer, vortexed 30 s and sonicated for 3 min at 130 W. After second round of 30 s vortex, samples were centrifuged at 6000 × g and 4°C for 10 min. The pellet was re-suspended in saline, OD_590nm_ was measured and appropriate dilutions were plated to assess adherent bacterial loads. Both, capsule and tissue were weighted and homogenized with saline. Appropriate dilutions were plated to evaluate the amount of bacteria per mg.

### Statistical analysis

All statistical analyses were performed using GraphPad Prism 6 software (GraphPad Software, Inc., La Jolla, CA USA). The following statistical tests were performed, otherwise stated on the figure legend. Paired ttest was applied for biofilm formation and mRNA quantification. Ratio paired ttest was used for eDNA quantification. For fibronectin binding assay and internalization experiment unpaired ttest with Welch’s correction was applied. For mouse catheter model of infection, Mann-Whitney test was used (unpaired nonparametric compared ranks).

## Aknowledgments

We would like to thanks Prof. Kenneth Bayles for his help and advices with the Nebraska Transposon Library, Dr Rosemarie Gaupp for the *rpiRc* mutant strains and Prof. Marc Chanson and Dr. Arnaud Lalive D’Epinay for their advices.

## Supporting informations

**Figure S1**. **Screening for biofilm regulators**. Each column represents one of the 108 mutant strain (symbols) compared to the wt (lines). Biofilm amounts (black triangles), DNaseI susceptibility (blue circles) and Dispersin B susceptibility (green squares) are represented vertically for each. Lines represent USA300 JE2 wt relative amount of biofilm (black line) and remaining biofilm after DNaseI (blue line) or DispersinB treatment (green line). *rpiRc* is shown with bigger sized symbols.

**Figure S2**. **Growth curves in TSBg.** Growth curves were performed in TSB supplemented with 1% glucose because the complementation strains were not able to grow in absence of glucose. All wt strains (blue lines) and *rpiRc* mutant strains (red lines) had similar growth curves. In TSBg, the complementation strain (green interrupted line) showed a shift in planktonic growth of several hours depending on the strain. In strain UAMS-1 (B) and SA564 (C), complementation delayed growth for 5 hours while it was delayed for 10 to 12 hours in USA300 JE2 (A) and SA113 (D) respectively. SA113 produces a lot of large clumps at the end of growth. Although strains are growing in presence of agitation, in presence of glucose SA113 wt produces large clumps responsible for the discontinuous growth curve observed at the end of stationary growth phase (D, blue line). *rsbU* deletion (orange line) in UAMS-1 also presented a small shift similar to the double *rsbU-rpiRc* mutant (interrupted purple line; E), while in SA564 *rsbU* single and double deletion (orange and interrupted purple lines) had similar planktonic growth curve compared to wt (F).

**Figure S3. SA113 biofilm index.** The complementation strain SA113 Δ*rpiRc* Nat7 has a slower planktonic growth. Biofilm index allows estimating the amount of sessile cells versus planktonic cells. This method does not take into account planktonic growth differences. Biofilm index was calculated as follows: Biofilm OD_595nm_ / (Biofilm OD_595nm_ + planktonic OD_595nm_). This method is based on previous biofilm index determination (8406839; 25255085; 14679235; 26659110). Biofilm OD595nm corresponds to the OD of CV staining. Planktonic OD595nm correspond to the OD of cells recovered from pooled biofilm and washing supernatants. Using this normalization, we could confirm that complementation do not resulted in decreased biofilm formation in strain SA113 after 6hours of biofilm formation. The decreased CV OD595nm observed in figure 3 is related to SA113 Δ*rpiRc* Nat7 slow growth only. This biofilm index also confirms the absence of differences between wt and *rpiRc* mutant strains for SA113.

**Figure S4. Triton induced autolysis.** Induced autolysis was tested as previously described (1), using triton X100 0.1%, and growth monitored for 5 hours on a TECAN reader. After addition of triton, bacterial OD595nm rapidly decreased for both USA300 JE2 wt and rpiRc mutant strain without any differences, meaning that increased eDNA production in the wt strain is most probably not related to *atl* mediated autolysis.

**Figure S5. Oxacillin induced lysis.** Murein hydrolase lysis was tested using oxacillin induced autolysis test following previously published method (1). First oxacillin MIC was determined for all the strains (0.5µg/mL, 0.25µg/mL and 64µg/mL for UAMS-1, SA564 and USA300 JE respectively). MICs were exactly the same for wt, Δ*rpiRc* and Nat7 strains. Oxacillin was added after 4h of growth. Growth in presence of 1x MIC, 2x MIC and 20x MIC was monitored (OD595nm on a TECAN reader) for additional 12 hours. Mean of 5 experiments with standard error of the mean is represented. Bacterial growth was normalized to the first OD595nm measurement (t0 = 100%). RpiRc deletion resulted in increased oxacillin induced autolysis in UAMS-1 starting at 2x MIC, while increased oxacillin induced autolysis starts at 1x MIC in SA564 strain. USA300 JE2 Δ*rpiRc* showed small decreased growth in presence of 1x and 2x MIC, but the growth profile is close to wt growth (similar slope). In addition, at 20x MIC, no differences could be monitored between wt and rpiRc mutant strain. All these strains are affected in their mature biofilm and eDNA amounts when rpiRc is deleted. The absence of conserved oxacillin induced lysis between these strains suggests that, although rpiRc could be involved in increased murein hydrolase dependent eDNA release in UAMS-1 and SA564, this mechanism is not the main regulated by RpiRc to modify eDNA amounts in USA300 JE2 mature biofilms.

**Figure S6. Biofilm susceptibility to antibiotics.** 24h old biofilms were challenged with increasing antibiotic concentrations for another 24h in 96 well plates. Antibiotic susceptibility was represented as OD difference between t6h and t0 after antibiotic challenge (reference 100% = control biofilm without antibiotic). C) and E) were performed in duplicates, therefore no statistical analysis was applied.

We tested antibiotics targeting different pathways in bacterial metabolism or directed against bacterial components. While E-test showed similar MICs between wt and mutant strains, *rpiRc* deletion always showed decreased biofilm recalcitrance to all these antibiotics.

**Figure S7. eDNA quantification in double mutants rpiRc-nuc1 and rpiRc-nuc2 in the biofilms of strain SA564.** Both extracellular nucleases have been deleted in strain SA564 wt and ΔrpiRc to determine whether these nucleases are involved in RpiRc mediated eDNA regulation. Single nuc1 and nuc2 mutants were also tested for eDNA content and we could not see any differences in eDNA amount compared to wt strain at 6h or 24h. Deletion of nucleases should increase the amount of eDNA in the biofilm (2). Mutations were confirmed by sequencing and growth on selection medium. In addition, each mutation (nuc1; nuc2; rpiRc-nuc1 and rpiRc-nuc2) was isolated twice independently and similar eDNA quantities were measured. Thus we only showed one mutant clone for each condition. Deletion of either nuc1 or nuc2 in the rpiRc mutant clone did not restore the amount of wt eDNA amounts at 6h or 24h. If the absence of eDNA modification in the nuc1 and nuc2 mutants is specific for this strain or related to our culture conditions, it do not have any role in RpiRc eDNA regulation. We did not add statistical test on the figure because only two biological replicates were performed. However, two sided paired ttest was computed and results are presented in Table S4.

**Figure S8. Gel page of biofilm SN from wt and Δ*rpiRc* strains.** Supernatant from 24h old biofilm of USA300 JE2 wt and Δ*rpiRc* strains were collected three times independently. Proteins were TCA precipitated and the pellet resuspended in a 2x Laemmli buffer before gel page run. We are looking for a protein that is prevalent in the mutant supernatant compared to the wt. The high density band observed at 35KDa was the most interesting. Selected bands used for peptide analysis are indicated in the figure (red rectangles). Positive (from the ladder) and negative controls were used to reduce noise and nonspecific peptide identification.

**Figure S9. HEK293 profile on the MoxiFlow cytometer.** Non fluorescent HEK293 cells were measured alone (no bacteria) in the MoxiFlow cytometer. The gap at 10µm splits HEK293 cells from dust and background particles, justifying the HEK293 size limit for our experiments at 10µm. In addition, fluorescence units of HEK293 cells alone do not reach the 60 FU we choose as the limit for fluorescence detection of stained bacteria internalized in HEK293 cells.

**Figure S10. Sequence of USA300 JE2 Δ*rpiRc* Nat7.** Native *rpiRc* sequence with a RBS was cloned within the pRAB11 plasmid following method described in the M&M section of the manuscript.

**Table S1. Standard error of the mean for qRT-PCR**

**Table S2. Two sided paired ttest for qRT-PCR**

**Table S3. E-test results**

**Table S4. Two sided paired ttest for double mutants *rpiRc*-*nuc1* and *nuc2* eDNA quantification**

**Table S5. Proteins in 24 hours old biofilms of USA300 JE2 wt and Δ*rpiRc***

